# In-Silico analysis reveals lower transcription efficiency of C241T variant of SARS-CoV-2 with host replication factors MADP1 and HNRNP-1

**DOI:** 10.1101/2020.11.22.393009

**Authors:** Armi Chaudhari, Minal Chaudhari, Sapna Mahera, Zuber Saiyed, Neelam M. Nathani, Shantanu Shukla, Chirag Patel, Dhaval Patel, Madhvi Joshi, Chaitanya G. Joshi

## Abstract

Novel severe acute respiratory syndrome-coronavirus-2 (SARS-CoV-2) has claimed more than 1.5 million lives worldwide and counting. As per the GISAID database, the genomics of SARS-CoV2 is extensively studied with more than 500 genome submissions per day. Out of several hotspot mutations within the SARS-CoV-2 genome, researchers have focused a lot on missense variants but the least work is done on the UTRs. One of the most frequent 5’ UTR variants in the SARS-CoV-2 genome is the C241T with a global frequency of more than 0.9. In the present study, the effect of the C241T mutation has been studied with respect to change in RNA structure and its interaction with the host replication factors MADP1 Zinc finger CCHC-type and RNA-binding motif 1 (hnRNP1). The results obtained from molecular docking and molecular dynamics simulation indicated weaker interaction of C241T mutant stem loops with host transcription factor MADP1 indicating reduced replication efficiency. The results are also correlated with increased recovery rates and decreased death rates of global SARS-CoV-2 cases.

## INTRODUCTION

According to WHO, COVID19 pandemic has affected over 40 million individuals from over 218 countries, areas and territories and has cost 1.1 million lives (1). SARS-CoV-2 belongs to the *Coronaviridae* family, is a ssRNA virus that mostly causes respiratory infections. This family comprises the lethal viruses like SARS, MERS, and SARS-CoV-2.

During a viral infection, there is a critical game of host-pathogen interaction, wherein either wants to survive and hence both pathogen and host tend to develop mechanisms for their existence. A majority of viruses contain their genome replication through 5’cap or have some secondary structures of RNA forming internal ribosomal entry sites (IRES) to recruit host replication factors. For viral replication, the 5’ untranslated region (UTR) makes a substantial contribution by providing important gear to regulate various host replication initiation factors in the case of RNA containing viruses, the majority in picornaviruses (2). Another mechanism includes non-AUG (Methionine) initiation where a particular virus target eIF2 inhibits host cell protein synthesis and lastly, to overcome the stress of certain antigenic peptides, few viruses use CUG (Leucine) initiation mode (3). Some viruses use another variant for translation initiation like the shutoff mechanism, which is an eIF2 independent mechanism, where viruses use the formation of downstream loop helix after initiation codon for efficient translation of viral mRNA (4). These are major mechanisms by which viruses occupy the host factors to use them for their own replication and translation. To prevent infection of viruses, these targets can thus to be efficient candidates to shut down the viral translation machinery (5). SARS-CoV-2 is likely to perform its replication through a 5’ cap-dependent mechanism (6). In SARS-CoV-2, an important helicase protein Nsp13 might be involved in the 5’ cap-dependent mechanism of viral replication. Nsp13 has RNA50-NTPase activity mediated by the NTPase active site of the nsp13-associated helicase domain (7) (8). Host factors involved in coronavirus proliferation include Annexin A2, which regulates IBV (immune bronchitis virus) frameshifting (9), hnRNP1 which binds with high affinity at MHV (−) strand of the leader sequence of RNA, PAPBs binds at poly-A tract of 3’UTR of BCov, TBV and TEGV (10) (11) (12), MADP1 Zinc finger CCHC-type and RNA-binding motif 1 binds at stem-loops structures at 5’ UTR of SARS-CoV-2 and IBV (13). The involvement of these factors is proven by wet-lab experiments involving *in vitro* pull-down assay and gene expression studies (14) (15).

SARS-CoV-2 after entry in the human host through ACE2 receptors enters the lung epithelial cells and injects its RNA into the host cytoplasm (16) (17). At the initial stage of infection, full-length negative-sense RNA copies are generated, which further act as a template for the synthesis of positive-sense genomic RNA (18). During the discontinuous replication process of the viral genome, RNA stem-loop structures SL1, SL2, TLR-S play an important role by interacting with host RNA binding proteins like Zinc finger CCHC-type and RNA-binding motif 1 MADP1. This interaction plays important role in viral replication and translation (14) (19) (20) (13). Novel SARS-CoV-2 seems to have stem-loop structures in 5’ UTR sequence and the translation is occurring through a cap-dependent fashion (21) (22) (23). A majority of them are reported as SL1, SL2, SL3, SL4A, and SL5. (24), (25), (26). SL1 contains a bipartite sequence that is possibly involved in the fine-tuning of viral RNA interaction with host protein. SL2 contains a tremendously conserved loop sequence of 5’-CUUCU (N)-3’ among most of the coronavirus species which is likely to form a tetra loop-like structure (26) (27). SL3 is eminently conserved among the group of gamma and beta coronavirus. SL3 comprises TRS-L (transcription regulating sequence) 5’-UCUCAA-3’, which is exposed in the core region of SL3 and meant to be highly involved in virus replication and transcription (22). SL4 is one of the major hairpin loops containing domain among the coronaviruses; located downstream to the TRS-L sequence and prone to mutation. Even a single or two SNP in SL-4 can change the frame of the sequence and be prone to abolish viral replication (25) (24) (26).

The genome sequence of SARS-CoV-2 is around 29-31 kb and more than 151K whole genome sequences have been deposited in GISAID by researchers across the globe. Top variants in the SARS-CoV-2 genome include high-frequency variants including C241T, C1059T, C3037T, C14408T, A23403G, G25563T and G28883C from which C241T variant had a 99% frequency with 0.505 entropy by October, 2020 (28). Though lot of research is published pertaining to the missense variants of the genome, to the best of our knowledge, no detailed reports on effect of the 5’ UTR variant C241T on host factor binding are available (29).

In this article, molecular docking and MD simulation studies were performed using the wild type (T) and mutant (C) sequences from 5’UTR of SARS-CoV-2 with host factor MADP1 and hnRNP1. Further, the wild type and mutant RNA-Protein complexes were analyzed for their stability by deciphering H-bonds formations, binding energy using MMGBSA, dynamics cross-correlation matrix and Principal Component Analysis.

## MATERIAL AND METHODS

### RNA secondary structure prediction

FASTA format sequence of 5’ UTR of SARS-CoV-2 from RFAM database with id RF03117 was used for the prediction of the secondary structure of wild type sequence and mutant sequence was generated using the same sequence by replacing C241 with T. RNA secondary and tertiary structure prediction is a very crucial thing to perform during *in silico* experiments to reach accurate conclusions. Strains of various SARS CoV-2 already have reported a presence of various stem-loops structure and bulges which is addressed in RFAM database; we took FASTA format sequence (around 300 base pairs) from RFAM id RF03117 for studying 5’ UTR sequences of beta coronaviruses (30).

RNA secondary and ternary structures were generated in RNA composer that uses the RNA FRABASE dictionary with the CHARM force field for generating effective minimized energy RNA structures (31). One plus advantage of RNA composers is the ease of predicting the ternary structures of long RNA sequences, which is a limitation of many RNA structure prediction software. The effect of single nucleotide variation in base-pairing probabilities was checked in mutaRNA (32). Final confirmation of folded RNA structure was confirmed in the X3DNA-DSSR Linux-based operating system which gives accurate root mean square deviation (RMSD) between predicted and experimentally verified (33).

### Prediction of RNA-protein binding sites

MADP1 binds to SL1 of SARS-cov2, which is selected as a protein binding site for the 5’UTR region. It is also known that TRS-L is involved in binding with hnRNP1 sequence 5’ CGGCUGC 3’, which was selected as a protein binding sequence (14) (23). Prediction of RNA residues binding to protein was also done by RNApin web server (34), where these sites are also falling in the protein binding region.

### Docking of protein-RNA complexes

In HADDOCK, molecular docking of host factors and RNA was performed (35) (36). Haddock is a popular docking program that performs docking using a data-driven approach. Docking encompasses three stages: rigid-body docking, semi-flexible refinement, and water refinement. In stage one for the docking of protein and RNA in their bound conformation, a total of 1000 structures were generated. Systematic sampling of 180 rotated solutions was used in the rigid-body docking stage to minimize the occurrence of false positives. The best 20% structures generated from the rigid-body docking stage were used in the second stage of semi-flexible refinement. The second stage further performs three-step refinement, rigid body torsion angle dynamics, and simulated annealing stage at different MD steps. In the final simulated annealing stage where 1000MD steps were performed from 300K to 50K with 2 fs time steps. In the final stage, both the side chain and backbone of protein residues at interface and RNA can move except the terminal base of RNA. This final stage consists of a gentle refinement (100 MD heating steps at 100, 200, and 300 K followed by 750 sampling steps at 300 K and 500 MD cooling steps at 300, 200, and 100 K all with 2 fs time steps) in an 8 Å shell of TIP3P water molecules (36). Docking of protein-RNA complexes were also performed in various servers like PATCHDOCK (37) (Schneidman-duhovny, Inbar, Nussinov, & Wolfson, 2005), NPDOCK (38), and HDOCK (39) to compare the docking score between two structures.

### Molecular dynamics simulation

Molecular dynamics simulation of Protein-RNA complexes was performed in Desmond implemented in Schrodinger (40) (41). The selected protein-RNA complexes were first immersed into a TIP3 water box, extending 10 Å beyond any of the complex’s atoms. Counter ions (Na^+^, and Cl^−^ ions) were added to neutralize charges. Salt concentration was set to 0.15 M sodium, and chloride ions to approximate physiological condition. The MD was performed in the NPT ensemble at a temperature of 300 K and 1.63 bar pressure over 20 ns with recording intervals of 1.2 ps for energy and 20 ps for trajectory. Simulations were run with the OPLS-3e force field compatible for Protein and nucleic acids. Plots and figures were generated in Pymol DeLano, W. L. 2009.

### Binding energy for protein RNA complex is calculated using the following equations

Binding energy components for four protein RNA complexes were calculated using PRIME module in Schrodinger using thermal_mmgbsa.py and residue-wise decomposition using breakdown_mmgbsa_by_resdiue.py script. The binding energy was calculated for a total of 1000 frames of the MD trajectory starting from the 1th ns until the end of the trajectory at the 100th ns. MMGBSA was calculated using the following equation.

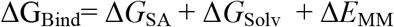

Δ*G*_SA_ is the difference in the surface area energies for the protein – RNA complex and the sum of the surface area energies in the protein and RNA. Δ*E*_MM_ is the difference in the minimized energies between the protein–RNA complexes. Δ*G*_Solv_ is the difference in the GBSA solvation energy of the protein–RNA complex and the sum of the solvation energies for the protein and RNA. Δ*G*_SA_ is the difference in the surface area energies for the complex and sum of the surface area energies in the protein and RNA (15).

### Principal component analysis

A vivid graphical presentation of dominant correlated motions of atoms present in the protein-RNA complex is obtained by the covariance matrix of the Cartesian coordinate data set. To generate covariance matrix of elements C_ij_ for coordinates i and j is given by bellow mentioned formula.

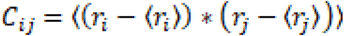

Here, r_i_ and r_j_ were Cartesian coordinates of i^th^ and j^th^ Cα atoms, 〈r_i_〉 and 〈r_j_〉 stood for the time average over all the configurations derived from the molecular dynamics simulation. This analysis was performed using the Bio3D library as implemented in R (42) (43) (44) (45).

### Dynamics cross-correlation

Cross-correlation Matrix was used to study detailed insights regarding to effect of C241T mutation on Protein-RNA complex dynamics by analyzing how atomic displacements were coupled. A cross-correlation coefficient C_ij_ was calculated from C_α_ atoms by the following equation.

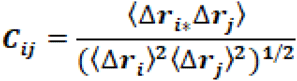

In this equation, Δr_i_ and Δr_j_ are the displacements from the mean position of the i^th^ and j^th^ residues (or atoms), respectively. The angular brackets “〈 〉” represents the time average of the entire trajectory. The value of C_ij_ is from −1 to 1. The positive value means positively correlated motion (moving in the same direction) indicated in cyan and the negative value represents negatively correlated motion (moving in the opposite direction) indicated in magenta. The DCCM analysis is carried out using the Bio3D packages of R (42) (43) (44) (45).

## RESULTS & DISCUSSION

Viral replication is highly dependent on the host factors, especially the transcription factors in the case of RNA viruses. Novel SARS-CoV-2, a positive-strand RNA virus from the family *Coronaviridae* is deploying 5’ CAP-dependent discontinuous replication within the human host (24, 46). As indicated in the previous studies in SARS infection, MADP1 from the Zinc finger protein motif 1 plays a very crucial role in deciding the replication efficiency within host cells (13). From hnRNP family, hnRNPA-1 is highly studied concerning SARS-CoV-2, having multiple roles in viral proliferation like RNA replication, transcription, export, import, and translation. hnRPA-1 uses RGG (Arginine-glycine-glycine) for RNA binding; it also binds with viral nucleocapsid protein (N) (47). While hnRNP1 is binding to the TLRS sequence (Pyrimidine rich) of most the coronaviruses like TGEV CoV and MHV (23), its involvement with respect to SARS CoV is yet be studied. In the present study, molecular dynamics and interaction of two human transcription factors MADP1 and hnRNP1 were studied with a most common 5’ UTR mutation C241T in the viral genome. MADP1 and hnRNP1 shares highly structural similarities, where an extra beta-sheet is found in hnRNP1 (Supplementary Figure S1). Due to similarities in tertiary structure and role of hnRNP1 in binding at TLR-s of other coronaviruses leads us to incorporate this protein in our study with respect to TLR sequence.

### Structural difference in 5’UTR of SARS-CoV-2

To initiate the same, sequences of the wild type and mutant (C241T) viral 5’ UTR sequences were taken and secondary structures were generated (Supplementary Table S1). RNA secondary structure which is predicted in dot-bracket format in X3-DNA Linux-based software (33) was visualized in Varna-GUI and structural differences due to C241T were interpreted. Representation of the variation in RNA structure due to SNP is elucidated in Table 1. The topological difference in both RNA structures is shown in Supplementary Figure S2. While structural features of RNA stem-loops are described in Figure 1. In mutant RNA, change is observed in SL-4 (101G-111U to 101G-112), which further leads to a change in the loop structure of SL4 (Figure 1). Change in SL4 also changes the folding; further inducing difference in SL1, SL2, and SL3. RNA fold was used to obtain nucleotide residue wise entropy and base pairing probability within residues (Supplementary Figure S3). Both RNA sequences showed structural and folding differences within SL1, SL2, and TLR sequences. To correlate these variations in sequence with the favorable binding of host replication factors, Protein-RNA docking and binding energy was calculated to check the effect of binding between both sequences of wild-type and (C241T) RNA. Studies have also shown the presence of these stem-loop structures in SARS, hence the resultant structures were taken to perform docking and MD simulation analysis (41).

**Table 1:**
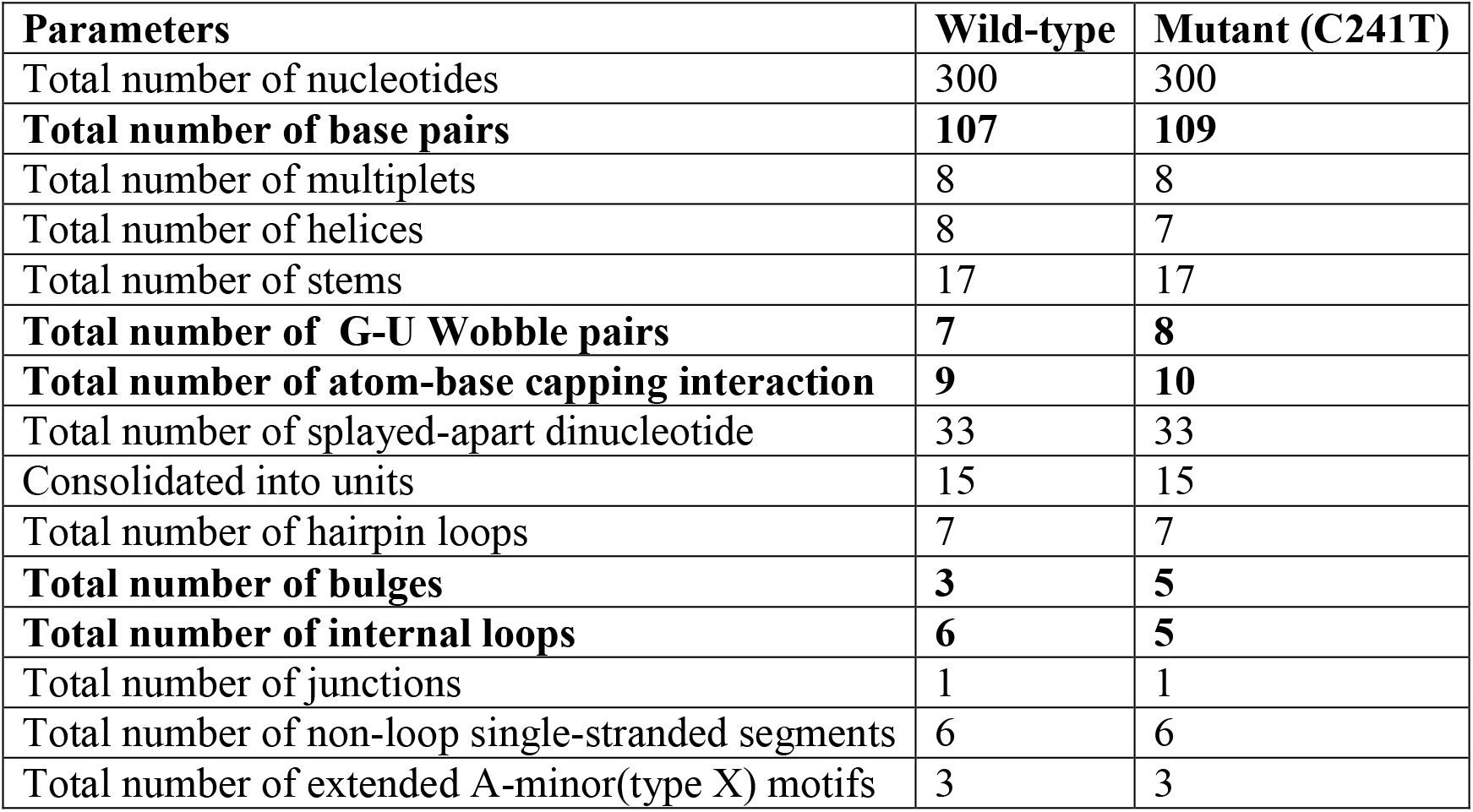
Tabular Representation showing the output of X3-DNA-DSSR in context of structural difference in both RNA sequences. Wild-type RNA sequences have changed in a number of base pairs, bulges, internal loops, and atom-based capping function compare to mutant. Change in below-mentioned parameters induces a change in the folding of SL1, SL2, and SL3 (shown in figure 1).

**Figure 1:**
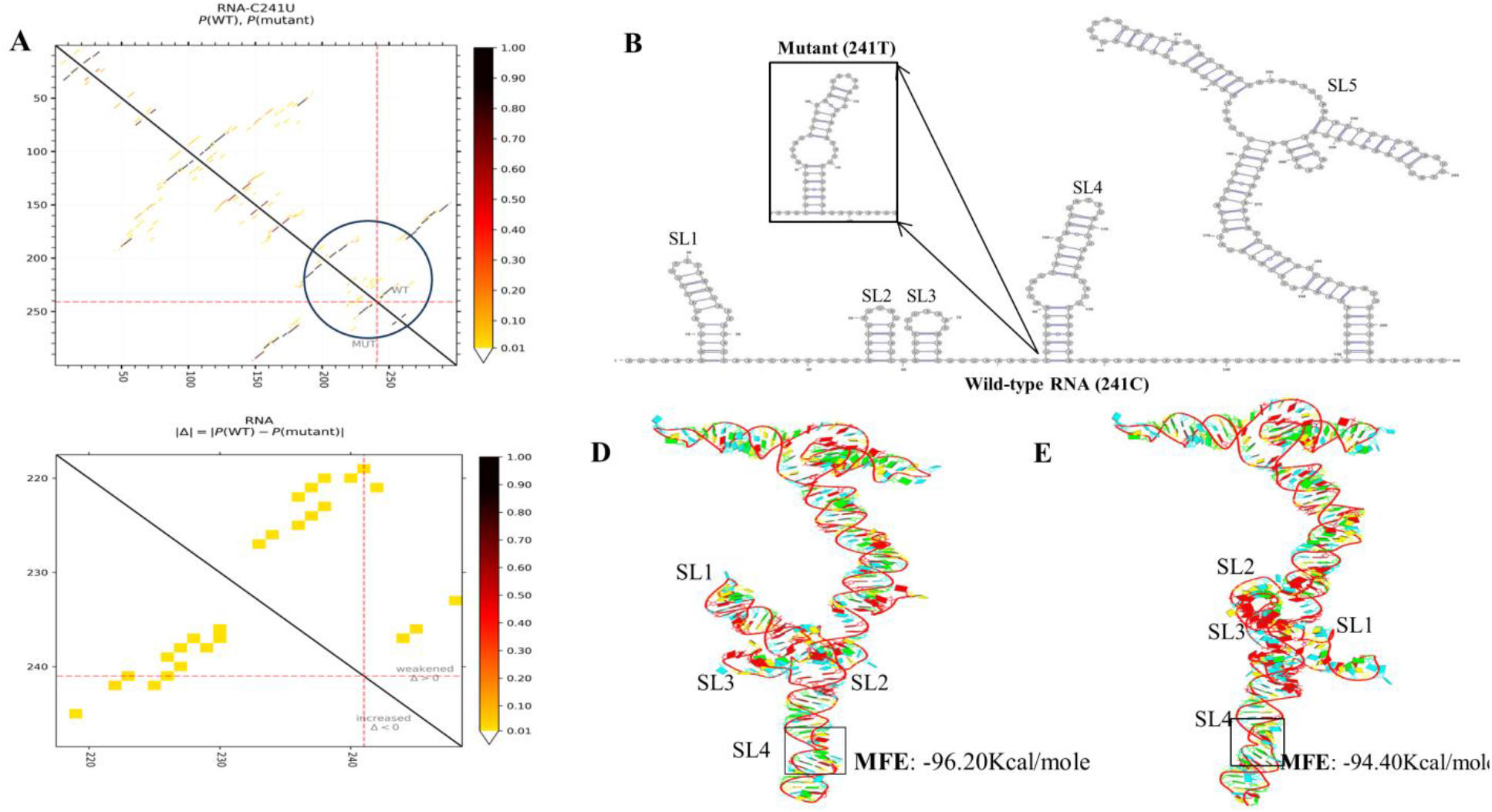
A.The base-pair probabilities for the wild type (WT) and mutant (MT) variant C241U of 5’ UTR of SARS-CoV-2. Its sequence context is plotted in the top-right and bottom-left halves of the matrix, respectively. The interval between two ticks on the axes represents 10 nucleotides. The evaluated mutation at position C241U is highlighted by the red lines along both axes. **B.** RNA 2D structure generated in efficient X3DNA-SSR Linux based system, to evaluated hairpin loops, bulges, stem-loops, multiples, no of wobble base pairs, extended A-minor(type X) motifs, and many more. Difference due to SNP C241U, C140U, and C292U is elucidated in **Table 1**. **C.** The differential heat map showing the base pair probability difference between the WT (Wuhan1) and mutant (C241U). A dark red (orange) color indicates a decrease (increase) in the pairing potential induced by the mutation. **D.** Main view is generated from X3-DSSR for wild-type RNA where SL1 and SL3 share nearby geometry. **E.** Main view generated from X3-DSSR for Mutant RNA where SL1 and SL3 share far from each other geometry.

### Docking of host transcription factors with 5’UTR variants

MADP1 is known to bind at SL1 and SL2 of RNA with help of Nsp1 protein for viral replication and transcription (13). Another protein hnRNP1 is known to bind at the TLR-S sequence of IBV, TGEV, and MHV (23). Wet lab reports of MADP1 binding to 5’ UTR of SARS-CoV were available but not for hnRNP1 (13). 5’UTR of SARS-Cov-2 is folded such that SL1, SL2, SL3, and SL4 shared nearby folding, which may enhance the surface area for binding of these proteins. Our major focus was on MADP1 mediated binding at the SL1 and SL2 regions of both variants of the 5’ UTR region. In the majority of *Coronaviridae* family viruses, hnRNP1 is known to bind at TLR-S sequence of 5’ UTR (11, 14). It was hypothesized that hnRNP1 might also bind to TLR-S of SARS-CoV-2 for viral proliferation. After docking studies, a change in binding was also observed for host factors with both RNA sequences. The stability of host factors-RNA complexes was also analyzed to comment on the affinity among both 5’ UTR variants.

Docking results for wild-type and mutant RNA with host factors protein from HADDOCK depicted the change in the folding of stem-loop 1, 2 and 3 in tertiary structures of RNA mediates difference in binding within all four complexes.

Concerning MADP1, a total of **117** structures in **8** cluster(s) were clustered in HADDOCK for wild type complex representing **58.5 %** of the water-refined models, while **119** structures in **8** cluster(s), were generated for the mutant sequence (C241T) which represents **59.5 %** of the water-refined models.

For HNRNP1, HADDOCK clustered **107** structures in **13** cluster(s), which represents **53.5 %** of the water-refined models generated for wild-type complex, while **92** structures in **14** cluster(s), which represents **46.0 %** of the water-refined models were generated for the mutant complex.

Based on the docking score, wild-type RNA seems to have better binding and RMSD value for MADP1 compared to the mutated (C241T) sequence (Table 2). The binding pose of MADP1 with RNA structure is shown in Figure 2. For hnRNP1, no significant change was observed for docking score and RMSD values. However, the average standard deviation (negative Z score) among clusters of wild-type RNA with host factors indicates better docking among complexes compared to the mutant (Table 2). Based on docking result, electrostatic energy and Van der Waals energy favors the protein-RNA complex formation while desolvation energy hinders the same.

**Table 2:**
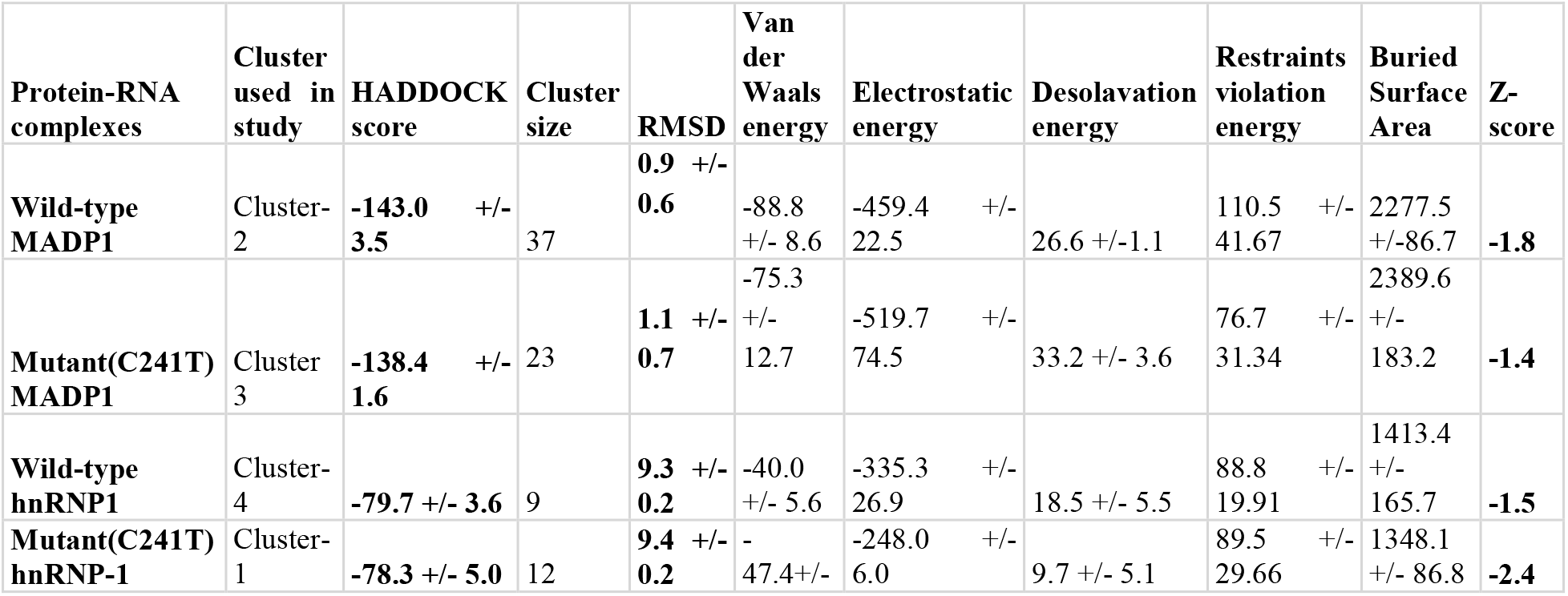
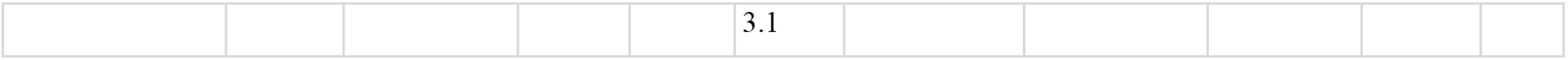
Docking results from HADDOCK. HADDOCK score for wild-type complex with respect to both host factors is more negative compare to the mutant. Parameters obtained from docking instigates that the wild-type complex seems to be more stable compare to the mutant in the case of both host factors, MADP1 and hnRNP1.

**Figure 2:**
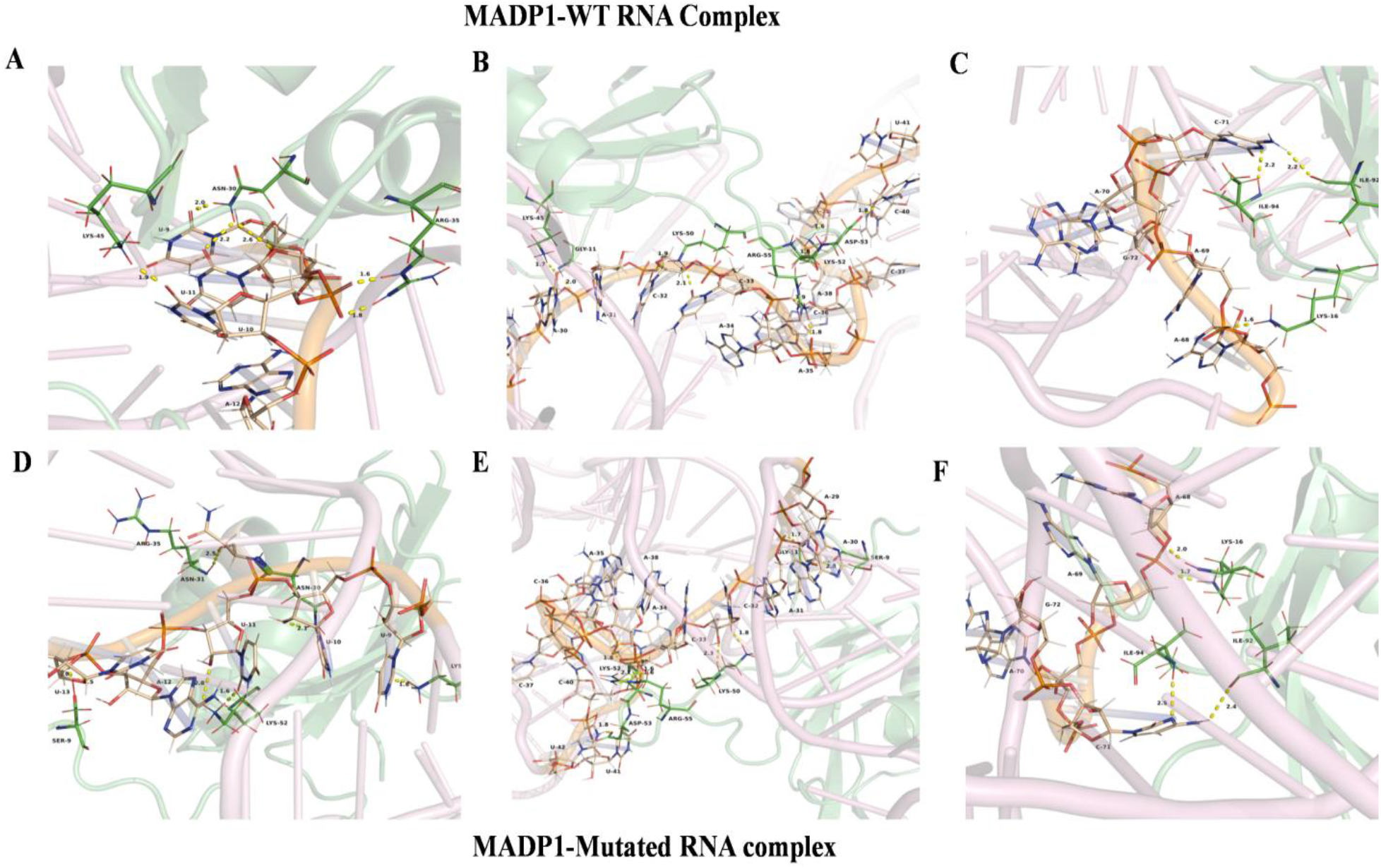
Docking pose generated from HADDOCK for MADP1-RNA complex. MADP1 seems to have interfacial hydrogen bonds and stacking interactions with RNA residues within SL1, SL2, and SL3 region of WT and mutant RNA.**2A & 2B**: MAPD1-WT RNA complex having pivotal interactions at SL1 region. **2C:** MAPD1-WT RNA complex having unique interactions in SL3.**2D& 2E**: MAPD1-Mutated RNA complex having pivotal interactions at SL1 region. **2F:** MAPD1-Muatated RNA complex having unique interactions in SL3.

**Figure 3:**
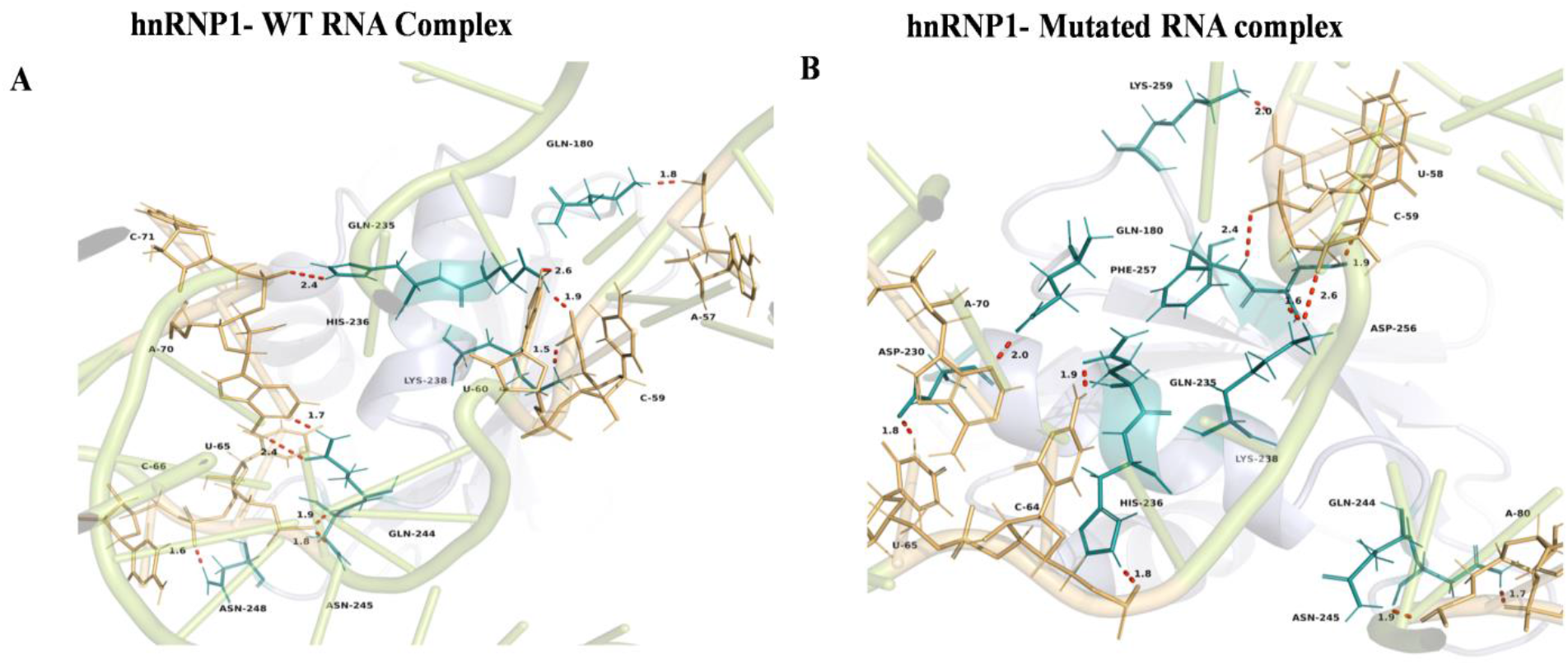
Docking poses generated from HADDOCK for hnRNP1-RNA complex. MADP1 seems to have interfacial hydrogen bonds and stacking interactions with RNA residues within the SL1 and SL3 (TLR-S) region of WT and mutant RNA.**3A**: hnRNP1-WT RNA complex having pivotal interactions at SL3 region containing TLR-S. **2C:** hnRNP1-Mutated RNA complex having interactions in SL3 region containing SL3.

As per the data analysis from GISAID and Nextstrain, the variant C241T emerged in early 2020 strains of SARS-CoV-2 and currently its most predominant variant in genomes reported from world-over (28, 48). To further validate this finding, the molecular dynamics of mutant and wild-type complexes were studied to find binding energy and stability.

### Analysis of MD simulation

Molecular dynamics simulation was performed for variants of 5’ UTR with two different host transcription factors MADP1 and hnRNP1 after docking in HADDOCK. Simulations of four complexes were performed for 100ns because early dissociation of MADP1 with Mutant (C241U) was observed (Supplementary Video S1 & S2). Early dissociation in mutant-MADP1 is one of the important factors to comment on the stability within both variants. Narrowing of space within SL1 and SL3 affects the binding MADP1 drastically which may lead to early dissociation. The major purpose of the Molecular dynamics simulation study is to comment on the stability within the 5’UTR variant and protein complexes. Various graphs captured during simulation events of both the wild type and variant in the simulation analysis package of Schrodinger are described in Figures 4 & 5.

**Figure 4:**
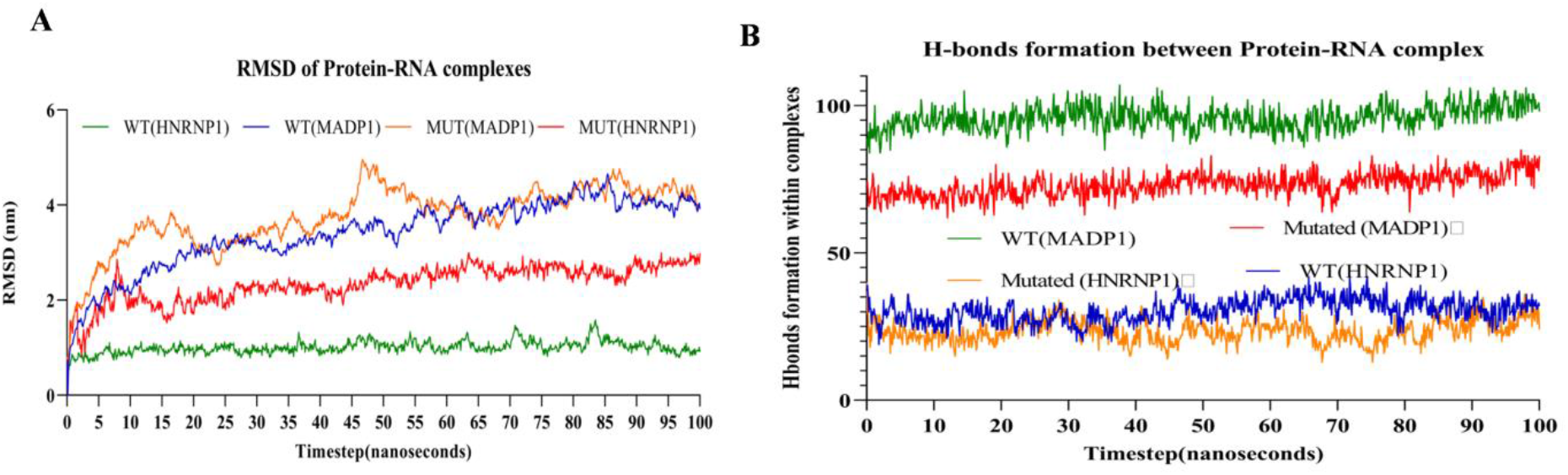
Analysis of MD simulation: RMSD and Hydrogen bond formation within complexes. **4A:** Wild-type complexes with respect to MADP1 and hnRNP1 shows lower RMSD values compare to mutant complex.**4B:** Hydrogen bond formation with respect to the wild-type complex is also high with respect to both proteins. Based on RMSD and H-bonds overall wild-type RNA shows efficient interaction compare to mutated RNA.

**Figure 5:**
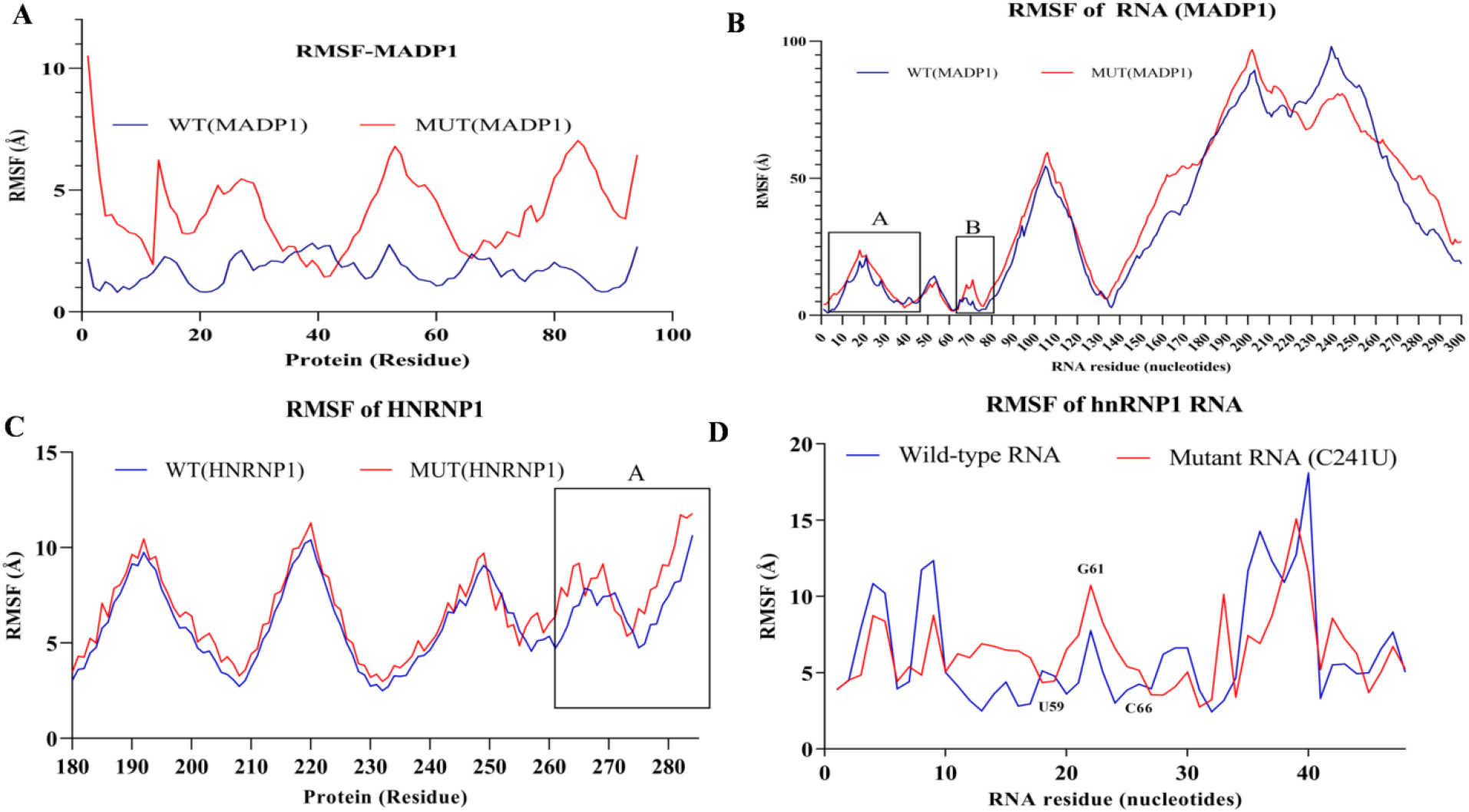
RMSF of protein and RNA residues within the complexes. **5A:** RMSF of MADP1 amino acid residues in wild type complex (blue) shows lower residual fluctuation compare to mutant complex (red). **5B:** Overall pattern of residual fluctuation in hnRNP1 is similar concerning both complexes, where Mutant complex’s hnRNP1 shows higher fluctuation compare to the mutant.**5C:** RMSF of in SL1 and SL2 region in MADP-RNA complex shows the lower fluctuation in wild-type complex, leading to efficient binding of MADP1 in wild-type RNA compare to mutant RNA. **5D:** RMSF of hnRNP1-RNA complex shows the lower fluctuation of TLR-S residues compares to mutant complex which is shown by labelling C59-C66 nucleotide residues.

With respect to MADP1, RMSD (Root mean square deviation) of wild-type complex was 5.37nm while for mutant it was 5.19nm. In the case of wild-type RNA, for the first 20ns trajectory was fluctuating but after 30ns system reached a plateau showing that the MADP1-RNA complex had formed a stable transition structure. While the trajectory of the mutant (C241U) complex was observed to be fluctuating and reaching a plateau after 60 ns (Figure 4A). RMSF (root mean square fluctuation) values of MADP1 and RNA from both complexes were calculated. Figure 5 explains the MADP1 residual fluctuation in both complexes, where amino acid residues from 1 to 60 show less fluctuation in wild type complex compared to the mutant (C241T) complex. Less fluctuation in RMSF value indicates more interaction between RNA-protein complexes (49, 50, 51) which is also evident here in the case of a wild-type complex.

HnRNP1 and RNA complexes were also analyzed with respect of stability. RMSD of the wild-type complex is 1.45nm which is much lower compared to mutant complex 2.35nm (Figure 4A). Overall residual fluctuation is shown in Figure 5C & 5D. Protein residues that are binding to RNA’s TLR-S sequence have overall less fluctuation compared to non-bound residues, among them wild-type hnRNP1 have lower fluctuation compared to mutant (Figure 5C-box A). TLR-S (U59-C66) of the wild-type complex has significantly lower fluctuation compare to the mutant complex (Figure 5D).

For both host replication factors, wild-type complex seems to be more favorable for the binding compare to the mutant complex. Further, Hydrogen bond formation was analyzed, where again in wild-type complex hydrogen bond formation is higher compared to mutant complex. Change in hydrogen bonds during different time intervals and occupancy within the complex was analyzed.

### Hydrogen bonds occupancy within complexes

To study the interaction of amino acids in Protein-RNA complexes, hydrogen bond analysis of those amino acid residues interacting with RNA was analyzed. Protein and RNA interaction with respect to hydrogen bond occupancy is shown in Supplementary Table S2. C241T mutation leads to structural changes in the folding of RNA (Supplementary Figure S2) which in turn affects the binding of MADP1 with SL1 sequence of 5’UTR and hnRNP1 with TLR sequence within SL3. H-bonds occupancy within 3Å distance cutoff and 20° distance was calculated using VMD (52). Based on docking results and MD simulation, hydrogen bonding was divided in two parts. Group 1 (named SL1) captured nucleotide resides ranging from 7G to C42 (Supplementary Figures S4 – S7), Group 2 (named TLR-S) contained 60U to 79A (Supplementary Figures S4 – S7). Hydrogen bond occupancy is shown for group 1 and group 2 in Tables 2 & 3, as those residues are exclusively involved in viral RNA replication using MADP1 and hnRNP1, respectively. RNA nucleotide residues involved in hydrogen bonding with amino-acid residues of MADP1 were shown using Pymol during different time intervals (0ns, 30ns, 60ns, and 100ns) of MD simulation. For group 1 (SL1) and group 2 (TLR-S), it is visible that the wild-type protein RNA complex is having more hydrogen bonds interaction with greater occupancy than the mutant (C241T) complex.

**Table 3:** Binding energy components from protein-RNA complex.

| **Energy components** | **WT**<br>**MADP1** | **C241U**<br>**MADP1** | **WT**<br>**hnRNP1** | **C241U**<br>**hnRNP1** |
| --- | --- | --- | --- | --- |
| $\Delta G$ Binding | -65.5821 | -54.6634 | -69.2351 | -45.8545 |
| $\Delta G$ Electrostatic energy | **-830.265** | **-188.509** | **-1253** | **-1049.28** |
| $\Delta G$ Covalent energy | 5.812749 | 2.775439 | 3.256262 | 11.40654 |
| $\Delta G$ Hbond energy | **-10.2371** | **-6.53** | **-13.99** | **-10.1914** |
| $\Delta G$ Lipophilic energy | -4.05164 | -4.36723 | -2.8212 | -4.96648 |
| $\Delta G$ pi pi interection energy | -9.59606 | -3.17418 | 1.616874 | -0.23087 |
| $\Delta G$ self contact correlation | -0.06446 | -0.00658 | -0.02245 | -0.17093 |
| $\Delta G$ Solv_GB | 861.3016 | 193.0531 | 1288.897 | 1101.954 |
| $\Delta G$ vdw energy | **-94.3725** | **-93.1687** | **-47.9052** | **-78.4821** |

Change in the tertiary structure of RNA leads to a drastic change in the binding of amino-acid residues for both of the proteins. Widening of a gap within the Wild-type complex leads to the binding of diverse MADP1 amino-acid residues with RNA, enhancing the stability compared to only one amino acid binding to two nucleotides. For example, in case of MADP1, a SL1 residue C32 is binding to Lys50-NZ in wild type complex while in mutant (C241T) Lys50 is binding with U9 and C32. U9 of wild-type complex is binding with Asn30-ND2, for mutant complex U10 is binding with the same amino acid (Supplementary Figure S4B & S4E). Occupancy of hydrogen bonds also increases in such type of interaction which is correlated with percentage occupancy results of Hydrogen bonds with MADP1 using VMD and decreased in bond length.

Some hydrogen bonds existed for the only period of time, named dynamic hydrogen bonds, while those that existed throughout the simulation are named as static hydrogen bonds. In group 1 there is a major change in interacting amino acids within SL1 region. Although, overall amino acid residues showed interactions were the same but change in position of interaction was observed, which is favoring higher occupancy of hydrogen bonds within wild-type complexes. Other interactions were also observed in wild-type complex, forming higher occupancy hydrogen bonds with amino acid residue in Lys-50, Asp-53, and Arg55 with 39A, 42U, and 33C respectively forming static hydrogen bonds (Supplementary Figure S5B & S5E, S6B & S6E). Lys-52 is the major amino acid favoring complex formation within both complexes. In wild type complex, Lys-52 was having bonds with C33, C36, and C38 throughout the simulation. In the mutant, Lys-52 was having bonds with C33 and 34A throughout the trajectory. Arg-48 seems to have bonds with 3A in the wild-type complex while again C33 in the mutant complex. Wild-type complex seems to cover wild range of nucleotides with different amino acids, which might make it more prone to efficient protein RNA complex formation. Vast varieties of dynamics of hydrogen bonds formed during the simulation are shown in the Supplementary Material.

Interesting results were observed when MADP1 is showing binding within TLR sequence in both complexes. In the 3D folded structure of 5’UTR, TLR-S residues share a nearby position with respect to SL1 and SL2. This unique interaction leads to one of the parameters to yet study, weather MADP1 is interacting in SARS-CoV-2 TLR-S sequence for viral proliferation or another factor is required for the same. At 100ns major change involved is in the mutant complex where protein is getting dissociated from RNA, compared to wild type complex (Supplementary Video S1 & S2). This can lead to the conclusion that the mutant complex is having relatively less stable interaction with host factor MADP1 compared to the wild type.

Other hypothesized protein-RNA complex (hnRNP1) is analyzed for commenting on the structural stability of RNA with respect to protein binding and interaction. It was surprising for us that Hydrogen bond occupancy in wild-type complex is much higher compared to the Mutant complex (Supplementary Table S3) shown in supplementary figures 8 and 9. Here also the same thing is repeated with respect to MADP1, where numbers of nucleotide residues binding to HNRNP1 were more in wild-type complex compared to the mutant. Nucleotide residues 57A, 58U, 59C, 60U, 64C, 65U, 69A, 70A were binding in wild type complex while 57A, 58U, 64C, 65U only binding in the mutant complex (Supplementary figure S8 & S9). However, no dissociation among both HNRNP1 and RNA was observed at 100ns.

RMSD, RMSF, H-Bonds and binding energy in form of MMGBSA indicated weak interaction of mutant protein-RNA complex than wild-type. Results were further correlated with DCCM and binding energy.

### Dynamics cross-correlation

Hydrogen bond interactions are correlated with dynamics cross-correlation matrix, where the interaction between protein and RNA with respect to four complexes is calculated throughout the trajectory of 100ns.

As shown in Figure 6, Dynamics cross-correlation matrix was calculated between inter and intra protein motions in both wild-type and mutant (C241T) complexes. As previously discussed in paper, SL1 and SL2 sequences are main interaction point in host-protein MADP1 for SARS-CoV-2-RNA during viral replication. DCCM of 1000 snapshots was performed. In DCCM matrix cyan colour indicates positive cross-correlation, magenta colour indicates negative cross-correlation, and white colour indicates no cross-correlation. DCCM was created for protein-protein, Protein-RNA and RNA-RNA interaction of Cα atoms in both complexes. DCCM for MADP1-RNA complexes were shown in Figure 6. SL1 and SL2 residues in wild-type complex are showing much higher cyan color intensity with respect to mutant complex (shown in box A). While in Box B nucleotide residues from SL3 were shown. These residues are TLR-S residues among 5’UTR sequence. MADP1 showing unique interaction within TLR-S sequence which is also depicting in DCCM (Box: B). Residues in Box A were showing positive correlation with C alpha atoms of protein at C_ij_ value of 0.25-0.85 in wild-type complex, while in mutant complex C_ij_ value was decreased to 0.00 to 0.125.RNA nucleotide residues no 59 to 75 shows positive cross-correlation with protein residues Lys-16, Phe57, Ile92, Ile94, Ala79, Lys-45, Thr-54, Arg-55, Arg35, Met-8, Ser-6, Gly-7, Pro14 and Lys68 in wild-type protein-RNA complexes with more frequency than mutant complex; which is depicted in box B in both figure 6A and 6B. Whereas for Box B , C_ij_ values for mutant complex (0.39) were lower than wild type complex (0.87). Wild-type complex’s RNA showed positive correlation in beta sheet residues (green colored) Lys52, Thr47, and Arg35 of MADP1, where this correlation was absent in mutant complex. Lys42, showed more positive correlation with wild-type RNA compared to Mutant RNA (C241T). From DCCM, it is evident that intensity of positive cross correlation in SL1-SL2 of wild type RNA is more compare to the mutant RNA with MADP1 (cyan).

**Figure 6:**
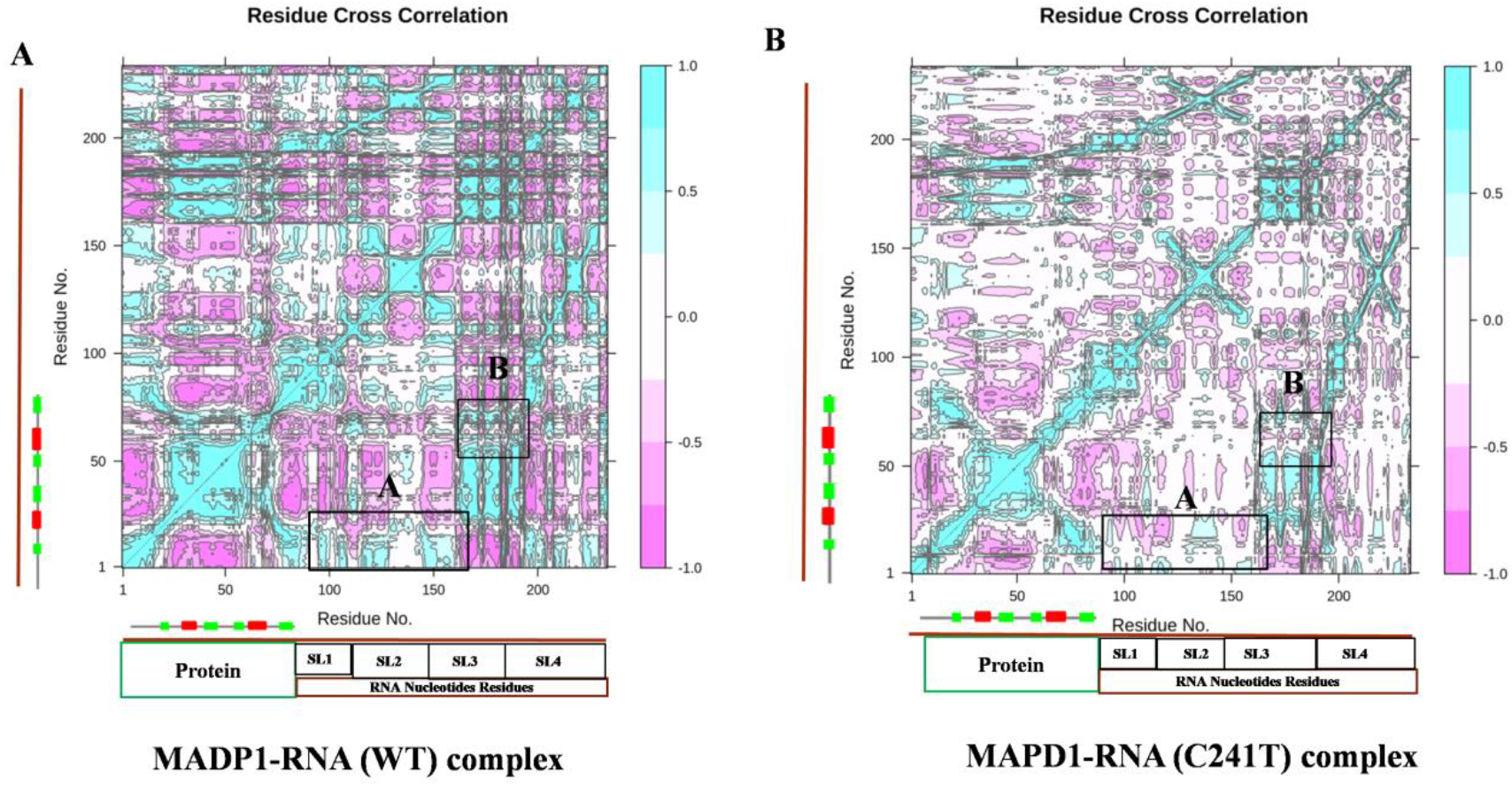
Dynamics cross-correlation for MADP1-RNA complexes. DCCM was calculated according to the time average of Cα atoms within the complex. The whole range of correlation from −1 to + 1 is represented in three ranges: cyan color corresponding to positive correlation values ranging from 0.25 to 1; magenta color corresponding to negative correlation values ranging from −0.25 to −1; and white color corresponding to weak or no-correlation values ranging from −0.25 to + 0.25. The extent of correlation or anti-correlation is indicated by variation in the intensity of respective cyan or magenta color.**6A**: DCCM of wild-type complex, box A and B shows a relatively higher frequency of cyan color compare to mutant complex in TLR-S sequence.**6B**: DCCM with respect to MADP1-mutant RNA complex.

DCCM of hnRNP1 is shown in Figure 7, where RNA nucleotide residues on C59, U60, G61, and U62 show interaction with protein amino acid residues Gln180, Lys283, Lys206, Gln253, Ile246 and Asn-48, Gln244 by the formation of hydrogen bonds within 3Å respectively. The intensity of TLR-S sequence is clearly visible that TLR sequence shows a strong positive correlation with wild-type RNA compared to the mutant which is shown in box A, B, C of both complexes (Figure 7).

**Figure 7:**
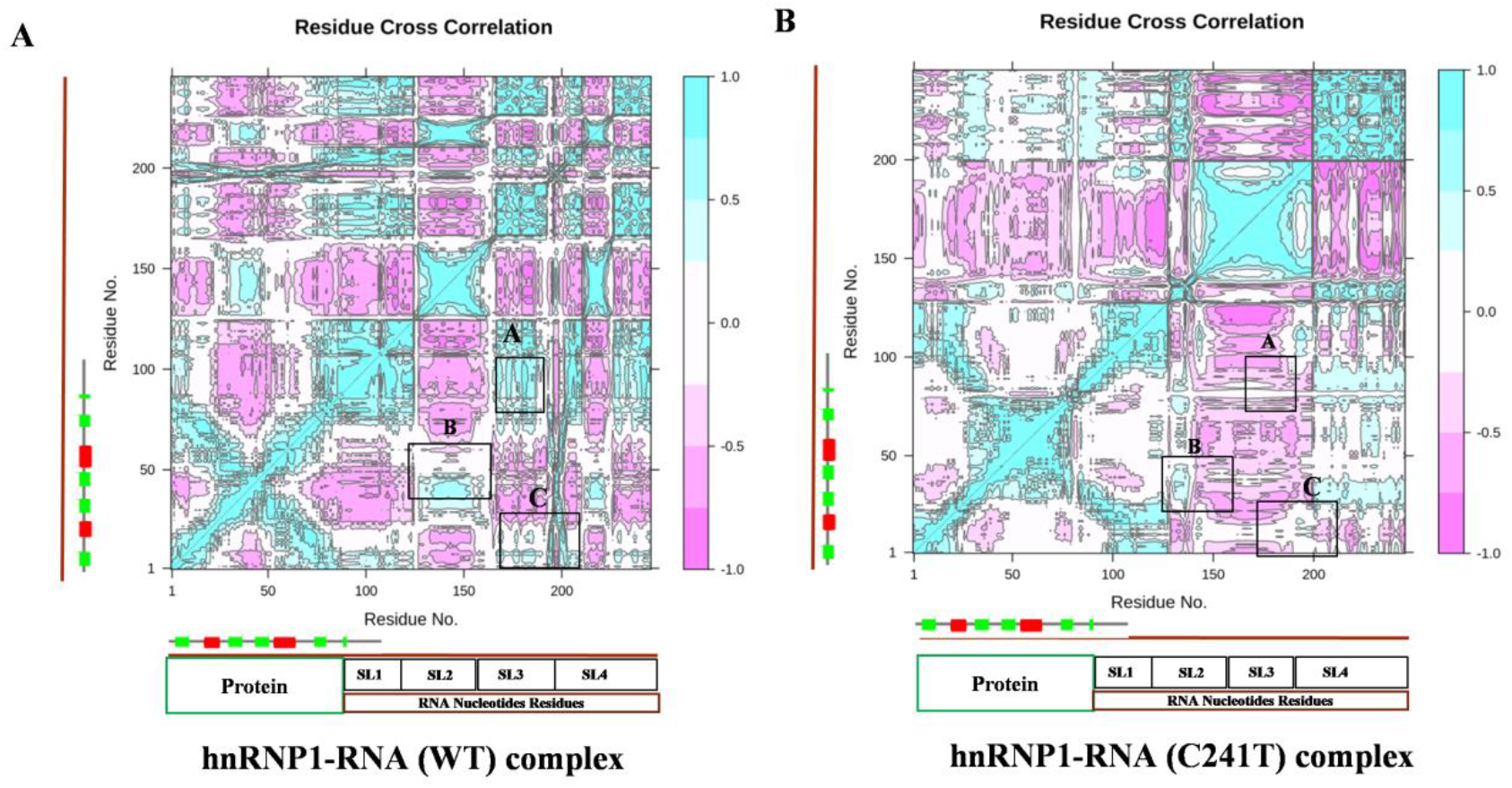
Dynamics cross-correlation for hnRNPN1-RNA complexes. DCCM was calculated according to the time average of Cα atoms within the complex. The whole range of correlation from −1 to + 1 is represented in three ranges: cyan color corresponding to positive correlation values ranging from 0.25 to 1; magenta color corresponding to negative correlation values ranging from −0.25 to −1; and white color corresponding to weak or no-correlation values ranging from −0.25 to + 0.25. The extent of correlation or anti-correlation is indicated by variation in the intensity of respective cyan or magenta color.**6A**: DCCM of wild-type complex, box A and B shows a relatively higher frequency of cyan color compare to mutant complex in the stem-loop region. **6B**: DCCM concerning hnRNP1-mutant RNA complex.

From DCCM it is clearly visible that residue interaction with host initiation factors shows strong positive correlation with nucleotide residue of wild-type complex compares to mutant complexes.

### Principle Component Analysis (PCA)

PCA was used to study the distinct protein conformational states in a principal component (PC) phase space during the MD simulations. The conformational transitions of the complexes were studied by projecting their trajectories onto a two-dimensional subspace spanned by the first three eigenvectors (PC1, PC2, and PC3) with respect to Cα residues. In Supplementary Figure S10 & S11, both complexes attained two conformational states on the subspace (shown in red and blue). The intermediate state located between these two conformations is shown with white dots.

For hnRNP1, PC1/PC2 and PC3/PC1 of wild-type shows thermodynamically distinct periodic jumps with a substantial energy barrier. While PC1/PC2 and PC2/PC1 of Mutant complex shows overlapping PC subspaces that lack energy barriers. Value PC1 in the wild-type complex is 71.95%, compare to 31.86% of PC1 in the mutant complex. The conjoined distributions of PC1/PC2, PC1/PC3, and PC2/PC3 of the Mutant complex shows energetically unstable complex (more scattered blue dots) compared to PC1/PC2, PC1/PC3, and PC2/PC3 of wild-type complex. More scattered system in the Mutant complex leads conclusion that, it is energetically less stable compared to wild-type.PC1/PC2 and PC3/PC1 of wild-type shows thermodynamically distinct periodic jumps with a substantial energy barrier. While PC1/PC2 and PC2/PC1 of Mutant complex shows overlapping PC subspaces that lack energy barriers (Supplementary Figure S11).

Same thing is observed in case of MADP1-RNA complexes, PC1 in wild-type complex is 64.35% and 73.42% of PC1 in mutant complex. More scattered system is observed PC2/PC3 suggests those principle modes are unstable in both complexes. PC3/PC1 of wild-type shows periodic jumps with sustainable energy barrier, in mutant scatterings suggests unstable system (Supplementary Figure S10).

Overall wild-type complex seems to be more stable compared to mutant (C241T) complex with respect to both proteins.

### Binding Energy components for Protein-RNA complexes

MMGBSA calculated for both protein-RNA complexes, where delta G bind is higher for wild-type complex compared to mutant complexes for both the proteins MADP1 and hnRNP1 (Table 3). First output structure containing minimized energy was taken in to study for both complexes. Residual decomposition was analysed to get the contribution of each residue within the complexes for favorable or unfavorable binding. To get insights regarding to energy components contributing more favourable binding of wild type complex, electrostatic, Hbonds, Van der Waals and ligand strain energy were analyzed.

For each protein-RNA complex, electrostatic energy is major energy component contributing to favourable binding within both complexes, while Solv GB (Generalized Born electrostatic solvation energy) significantly opposes the protein-RNA interaction and complex formation. Both are rather substantial in magnitude but compensate each other, overall resulting in favourable total electrostatic contributions.

Our major aim was to study the effect the mutation C241T with respect to host replication factor MADP1, docking and molecular simulation studies shows the effect of mutation in favour of host. Binding energy calculations are very crucial for commenting on the complexes formation concerning both wild-type and mutant RNA. For MADP1, binding energy for wild-type and mutant complex is −65.58kcal/mole and −54.66kcal/mole. From binding energy, wild-type complex seems to have more efficient complex formation compare to mutant. In figure 9 and 10, it is clearly visible that majority part of amino acid residues are falling in lime green-blue zone which represents near to zero value for energy contribution in mutant complex. While in the wild-type complex, residues favouring negative spontaneous energy are significantly high.

In wild-type complex, amino acid residues interacting more spontaneous are Ser-1, Ser-2, Gly-7, Ser-9, Met-8, Gly-11, Tyr-34, Met-49, Ile-48, Lys-43, Arg-55 and Lys-52, showing interfacial hydrogen bonds and intermolecular stacking interactions with nucleotide residues A6, U9, U11, U21, U19, G23, G24, and A35. Residues decomposition graph for MADP1 residues and WT-RNA shows higher negative energy compare to other basic residues within the complex. It is interesting to study MADP1 residues 1-11 in both complexes, where significant change was observed with respect to interaction. In wild-type complex those residues (Ser-1, Ser-2, Gly-7, Ser-9, Met-8, and Gly-11) are involved in direct contact with RNA nucleotide residues C19, C21, G23, and G24 shown in figure 9E. Direct interaction shows more compactness in structure, which is also correlating with RMSF graph of MADP1 (Figure: 5B). While in mutant complex from those amino acid residues (1-11), Gly-1, Ser-2, Ser-5, Leu-6, Ala-7, and Pro-8 shows positive decomposition energy (and no energy), and leads to more fluctuations, hence less compact structure Wild-type (Figure 5B). Although for mutant complex those residues posses’ high electrostatic and Van der Waals energy but more positive Solv-GB energy of those residues hinders the complex formation. In wild type complex MADP-1 residues Lys-52 (25.32kcal/mole), Met-49 (−15.08kcal/mole), Gly-7 (− 27.30kcal/mole), and Ile-48 (−20.17kcal/mole) shows 4-5-fold higher energy decomposition compared to other basic residues (Figure 8A & 8B). In mutant complex amino acid residues interacting with more spontaneous energy are Ser-38, Lys-39, Tyr-40, Asp-67, Ser-70, Val-46, Tyr-34, and Asn-31 (showing direct contact with nucleotide residues) (Figure 10E). From Figures 9 and 10, it is clear that the majority of amino acids residues where electrostatic potential, Vander Waals and H-bonds interaction are high also shows high mmgbsa energy. Amino acid residues Lys-39 (−21.30kcal/mole), Tyr-40 (−23.60kcal/mole), Ser-38 (− 5.63kcal/mole), Asn-31 (−15.63kcal/mole) and Tyr-34 (10.63kcal/mole) are interacting with nucleotide residues at SL1 region of RNA and shows higher energy decomposition compared to the other non-interacting residues. MADP1-WT complex have 11 amino acid residues binding to nucleotide residues of RNA, compare to 8 amino acid residues binding to the mutant RNA. Overall MADP1 shows a higher binding affinity to SL1 region in WT RNA compare to mutant RNA.

**Figure 8:**
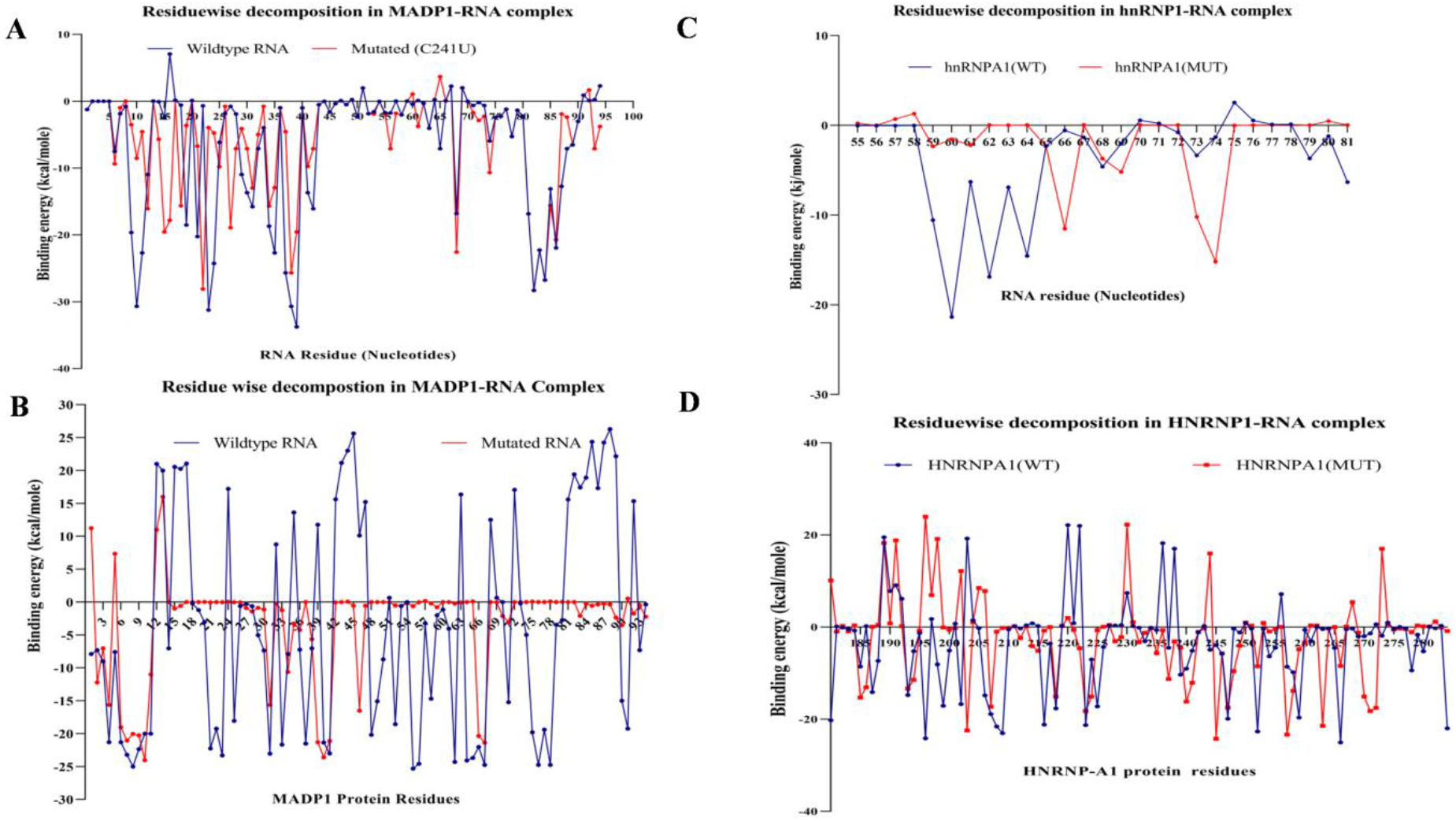
Residue wise decomposition obtained through MMGBSA for every four complexes. **8A**: Residue wise decomposition for RNA residues in MADP1-RNA wild-type and mutant complexes.**8B**: Residues wise decomposition of MADP1 in both complexes. **8C**: Residue wise decomposition of RNA residues in hnRNP1 RNA complexes. **8D**: Residues wise decomposition of hnRNP1 in both complexes

**Figure 9:**
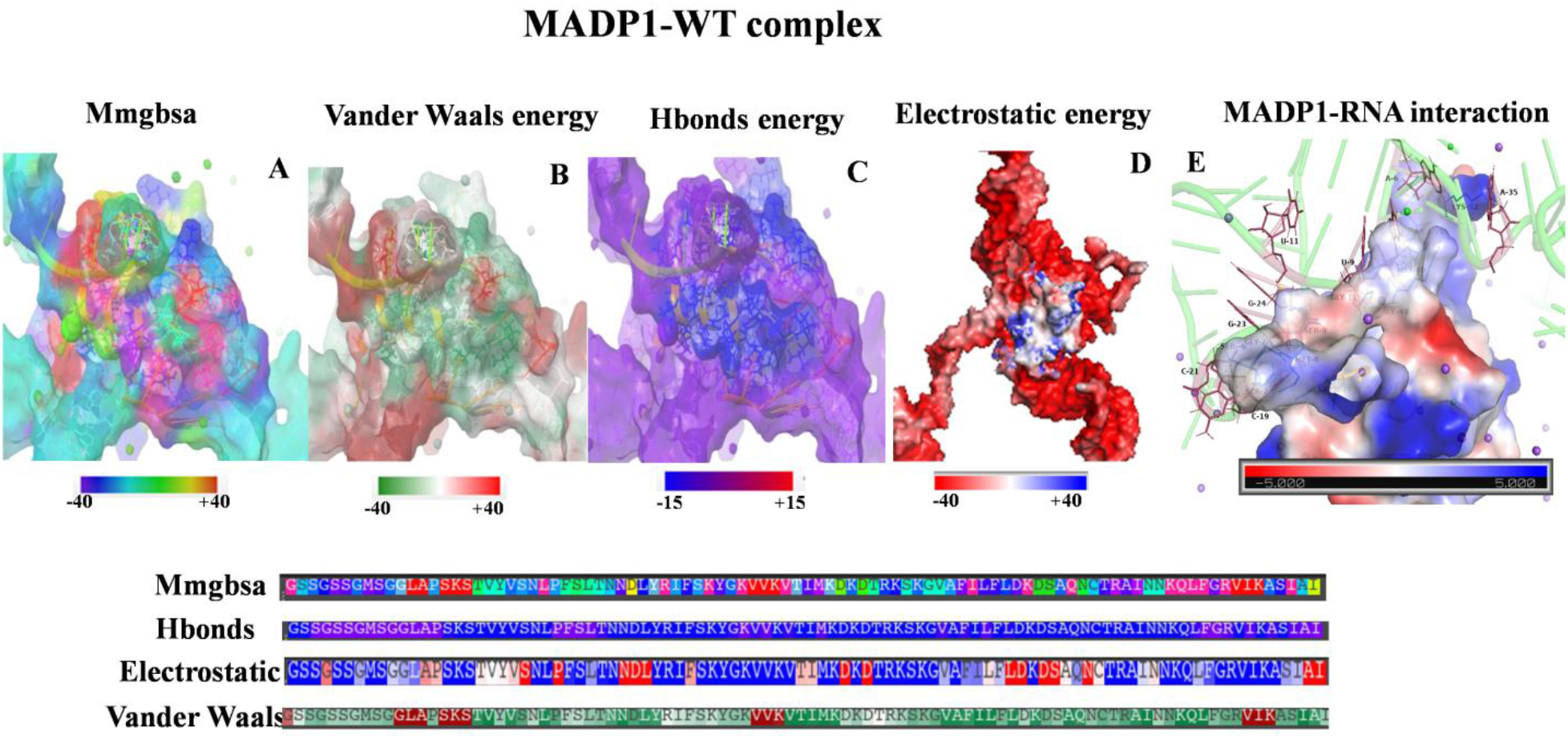
Contribution of Major energy components involved in the formation of MADP1-Wild-type RNA complex. Energy components contribution was done in the energy visualization package in maestro by entering cut-off values of −40 to +40 kcal/mole for MMGBSA, electrostatic potential, and Vander Waals energy. −10 to +10 cut-off values for Hydrogen bond energy.**9A:** MMGBSA for complex. **9B:** Vander Waals energy **9C:** Hbonds energy **9D:** Electrostatic energy and **9E:** MADP-WT RNA interaction in terms of electrostatic energy, as RNA was showing negative electrostatic potential due to the phosphate backbone it was not shown for a better image of interaction (for whole electrostatic interaction figure **9E** can be referred)

**Figure 10:**
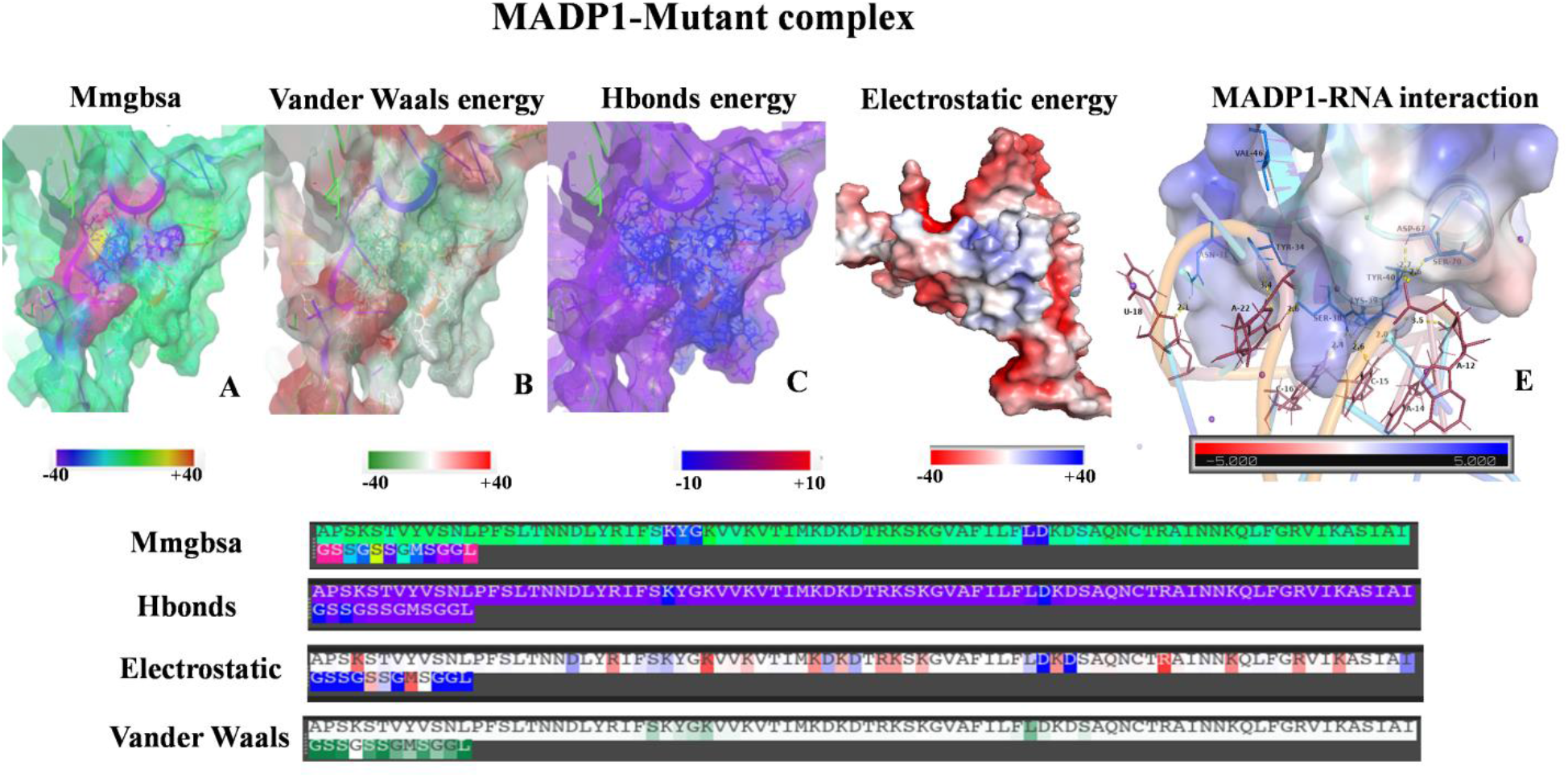
Contribution of Major energy components involved in the formation of MADP1-mutated RNA complex. Energy components contribution was done in the energy visualization package in maestro by entering cut off values of −40 to +40 kcal/mole for MMGBSA, electrostatic potential, and Vander Waals energy. −10 to +10 cut-off values for Hydrogen bond energy. **10A:** MMGBSA for the complex. **10B:** Vander Waals energy **10C:** Hbonds energy **10D:** Electrostatic energy and **10E:** MADP- mutated RNA interaction in terms of electrostatic energy, as RNA was showing negative electrostatic potential due to the phosphate backbone it was not shown for better image of interaction (for whole electrostatic interaction figure **10E** can be referred)

For hnRNP1, binding energy for wild-type and mutant complex is −69.85 Kcal/mole and −45.85 Kcal/mole. Again wild-type complex seems to bind target with higher affinity compare to mutant in case of hnRNP1 also. Other major contributing energy components for complexes are Vander Waals energy and hydrogen-bond energy (Table 3). These three energies are favoring complex formation more efficiently so energy potencies of each residue in complex were analysed. As electrostatic energy is favoring the complex formation, residues taking part in interaction are shown in term of electrostatic energy (Supplementary Figure S12D and S13D). With respect to protein residues, heat maps were shown to correlate amino acid residues with respect to their energy for complex formation. Protein interface residues binding to RNA are showing positive electrostatic potential complimentarily to negative electrostatic potential of the phosphate backbone of RNA (Supplementary Figure 12E and 13E). Extra positive electrostatic potential of residues, Lys 218, L, Lys-259, Arg-277, Arg-254 and Arg185 favors enhanced binding to TLR-S sequence of RNA, henceforth favouring efficient binding of wild-type complex (shown in blue) (Supplementary Figure 12E). To identify key residue important for binding, residual decomposition was analyzed for both complexes. For wild type complex RNA residues, U47, C59, U60, U62, U63, C64, A79, A81, C123 and G124 are interacting hnRNP1, covering TLR-S sequence -UCUU-, SL2 and SL4. In mutant complex RNA residues, U11, U10, C32, A31, C66, A69, A68, A73 and A74 seem to be interact with hnRNP1. It was quite surprising that the minimized structure of mutant was interacting within SL1 region, more efficiently compare to TLR-S region. HnRNP1 is supposed to bind at TLRs region for viral proliferation, mutated complex (energy minimized structure) is binding less efficiently to TLR-S region. Superimposition of both minimized structures gives the clear idea for the above statements, wt-hnRNP1 is near to SL3 having TLR-S and SL4, while mut-hnRNP1 is near to SL2 and SL3 (Supplementary Figure S12). From free energy decomposition, it was unsurprised that residues which were having more hydrogen bond energy and electrostatic energy will possess higher decomposition energy which correlating in figure 8. In wild-type complex RNA residues within the TLR-S sequence possess more spontaneous, decomposition energy U60 (21.36kcal/mole), U62 (16.89kcal/mole) and C64 (−14.56kcal/mole) compare to the average of −1.5-2 kcal/mole with same resides of the mutant complex. While in mutant residues, C66 (−11.53kcal/mole), A69 (−5.24kcaal/mole), A73 (−10.2kcal/mole), and A74 (−15.2kcal/mole) (Figure 8D). Wild-type sequence seems to have more favourable binding in TLR-s sequence compare to mutant, so 241C seems to be favourable for viral proliferation compare to 241U.

### Epidemiological data

Structural analysis of both RNA, reveal changes in TLR-S region while folding in 3D form, which is affecting the binding of host replication factor MADP1. According to results obtained through MD analysis Mutant, the complex seems to be less stable compare to wild type based on RMSD and Hydrogen bonding between protein RNA complexes. *In-Silico* findings suggest weaker interaction of mutant RNA with host factor MADP1 compared to wild-type. This suggests lesser replication efficiency in mutant SARS-CoV-2 strain. The 5’ UTR variant C241T has emerged in March, 2020 and its one of the most observed variants in genomes sequenced in 2020, with a frequency of 0.505 (22). To further correlate this finding, epidemiological data of global cases, death rate, and the recovery rate was obtained from World Health Organization covid-19 dashboard. The recovery rate in global cases has increased from 2.2% in March 2020 to 70% to date whereas the overall decline in death rate is observed (Supplementary Figure S15). Although there are many other reasons for increased recovery rate, but the effect of most frequent mutation in the genome can also be not neglected. According to WHO, before the ending of March death rate is 7.1% while recently in October the death rate is 1.46 %.

The fact that if viral RNA is binding less efficiently with host replication factors due to structural changes, the efficiency of virus replication within the host decreases. However, these findings can be further validated experimentally using cell lines and other *In-vitro* methods.

## Conclusion

5’ UTR interactions with host factors were studied and it was concluded that C241T changes the SL4, which overall changes the folding of RNA. MADP1 and HNRNP-1 reduce binding shows a decrease in efficiency of the virus to replicate in a host, which in terms decrease the death rate (increasing the recovery rate). Results were correlated with the epidemiological data which is also showing an increased recovery rate all over the world. HnRNP1 interaction which SARS-CoV two is not yet been studied. From our studies, it can be hypothesized that hnRNP1 is binding to the -UCUU sequence present in SARS-Cov-2.

## AVAILABILITY

1. Epidemiological statistics for SARS-CoV-2 was obtained from WHO (world health organization) link: https://covid19.who.int
2. https://trackthevirus.info/

## ACCESSION NUMBERS

Public repository accession numbers of Protein and RNA sequences used for this study are listed below.

1. Crystal structure of a Raver1 PRI3 peptide in complex with poly-pyrimidine tract binding protein RRM2 (**PDB: ID 3ZZY**).
2. Solution structure of RNA binding domain in Zinc finger CCHC-type and RNA binding motif 1, MADP1 (**PDB: ID 2E5H**).
3. Severe acute respiratory syndrome coronavirus 2 isolate Wuhan-Hu-1, complete genome **GenBank: MN908947.3.**

## SUPPLEMENTARY DATA

Supplementary Data are available at NAR online.

## ACKNOWLEDGEMENT

We are extremely grateful to the Schrodinger team for providing us an evaluation licence. We are also thankful to Dr Prajwal Nandedkar for providing constant help regarding to the Schrodinger software access and use. We also acknowledge Dr Saumya Patel for manuscript proofreading.

## REFERENCES

1. WHO,W.H.O. (2020) Situation Report - 65 - Coronavirus disease 2019. World Heal. Organ., 2019, 2633.

2. Martinez-Salas,E., Francisco-Velilla,R., Fernandez-Chamorro,J. and Embarek,A.M. (2018) Insights into structural and mechanistic features of viral IRES elements. Front. Microbiol., 8, 1–15.

3. Schwab,S.R., Shugart,J.A., Horng,T., Malarkannan,S. and Shastri,N. (2004) Unanticipated antigens: Translation initiation at CUG with leucine. PLoS Biol., 2.

4. Toribio,R. and Ventoso,I. (2010) Inhibition of host translation by virus infection in vivo. Proc. Natl. Acad. Sci. U. S. A., 107, 9837–9842.

5. Deng,J.X., Nie,X.J., Lei,Y.F., Ma,C.F., Xu,D.L., Li,B., Xu,Z.K. and Zhang,G.C. (2012) The highly conserved 5′ untranslated region as an effective target towards the inhibition of Enterovirus 71 replication by unmodified and appropriate 2′-modified siRNAs. J. Biomed. Sci., 19, 1.

6. k. nakagawa, k.g. lokugamage, s. makin. (2020) free information in English and Mandarin on the novel coronavirus COVID-Viral and Cellular mRNA Translation in Coronavirus-Infected Cells. Adv. viral Res., ISSN 0065-, 77.

7. Lal,S.K. (2010) Molecular biology of the SARS-coronavirus. Mol. Biol. SARS-Coronavirus, 10.1007/978-3-642-03683-5.

8. Ivanov,K.A., Thiel,V., Dobbe,J.C., Meer,Y. Van Der, Snijder,E.J. and Ziebuhr,J. (2004) Multiple Enzymatic Activities Associated with Severe Acute Respiratory Syndrome Coronavirus Helicase. 78, 5619–5632.

9. Kwak,H., Park,M.W. and Jeong,S. (2011) Annexin A2 binds RNA and reduces the frameshifting efficiency of Infectious Bronchitis Virus. PLoS One, 6.

10. Spagnolo,J.F. and Hogue,B.G. (2001) Requirement of the poly(A) tail in coronavirus genome replication. Adv. Exp. Med. Biol., 494, 467–474.

11. Galán,C., Sola,I., Nogales,A., Thomas,B., Akoulitchev,A., Enjuanes,L. and Almazán,F. (2009) Host cell proteins interacting with the 3′ end of TGEV coronavirus genome influence virus replication. Virology, 391, 304–314.

12. Wu,H.Y., Ke,T.Y., Liao,W.Y. and Chang,N.Y. (2013) Regulation of Coronaviral Poly(A) Tail Length during Infection. PLoS One, 8, 1–12.

13. Tan,Y.W., Hong,W. and Liu,D.X. (2012) Binding of the 50-untranslated region of coronavirus RNA to zinc finger CCHC-type and RNA-binding motif 1 enhances viral replication and transcription. Nucleic Acids Res., 40, 5065–5077.

14. Fung,T.S. and Liu,D.X. (2019) Human Coronavirus: Host-Pathogen Interaction.

15. Guan,B. (2010) TRACE: Tennessee Research and Creative Exchange 5’ -Proximal cis-Acting RNA Signals for Coronavirus Genome Replication.

16. Zhou,P., Yang,X. Lou, Wang,X.G., Hu,B., Zhang,L., Zhang,W., Si,H.R., Zhu,Y., Li,B., Huang,C.L., et al. (2020) A pneumonia outbreak associated with a new coronavirus of probable bat origin. Nature, 579, 270–273.

17. García,L.F. (2020) Immune Response, Inflammation, and the Clinical Spectrum of COVID-19. Front. Immunol., 11, 4–8.

18. V’kovski,P., Kratzel,A., Steiner,S., Stalder,H. and Thiel,V. (2020) Coronavirus biology and replication: implications for SARS-CoV-2. Nat. Rev. Microbiol., 10.1038/s41579-020-00468-6.

19. Castello,A., Fischer,B., Eichelbaum,K., Horos,R., Beckmann,B.M., Strein,C., Davey,N.E., Humphreys,D.T., Preiss,T., Steinmetz,L.M., et al. (2012) Insights into RNA Biology from an Atlas of Mammalian mRNA-Binding Proteins. Cell, 149, 1393–1406.

20. Abramo,J.M., Reynolds,A., Crisp,G.T., Weurlander,M., Söderberg,M., Scheja,M., Hult,H., Wernerson,A., Emacs,A., Distribution,U.E., et al. (2012) Individuality in music performance Springer International Publishing.

21. Chen,Y. and Guo,D. (2016) Molecular mechanisms of coronavirus RNA capping and methylation. 31, 3–11.

22. Rangan,R., Zheludev,I.N. and Das,R. (2020) RNA genome conservation and secondary structure in SARS-CoV-2 and SARS-related viruses: a first look. Rna, 10.1261/rna.076141.120.

23. Sola,I., Mateos-Gomez,P.A., Almazan,F., Zuñiga,S. and Enjuanes,L. (2011) RNA-RNA and RNA-protein interactions in coronavirus replication and transcription. RNA Biol., 8, 237–248.

24. Guan,B. (2010) 5’ -Proximal cis-Acting RNA Signals for Coronavirus Genome Replication.

25. MiguelRamos-Pascual (2019) Coronavirus SARS-CoV-2: Analysis of subgenomic mRNA transcription, 3CLpro and PL2pro protease cleavage sites and protein synthesis Corresponding autor: Miguel Ramos-Pascual.

26. Madhugiri,R., Fricke,M., Marz,M. and Ziebuhr,J. (2020) Since January 2020 Elsevier has created a COVID-19 resource centre with free information in English and Mandarin on the novel coronavirus COVID-19. The COVID-19 resource centre is hosted on Elsevier Connect , the company’s public news and information.

27. Yang,D., Liu,P., Giedroc,D.P. and Leibowitz,J. (2011) Mouse Hepatitis Virus Stem-Loop 4 Functions as a Spacer Element Required To Drive Subgenomic RNA Synthesis ◻. 85, 9199–9209.

28. Hadfield,J., Megill,C., Bell,S.M., Huddleston,J., Potter,B., Callender,C., Sagulenko,P., Bedford,T. and Neher,R.A. (2018) NextStrain: Real-time tracking of pathogen evolution. Bioinformatics, 34, 4121–4123.

29. Mercatelli,D. and Giorgi,F.M. (2020) Geographic and Genomic Distribution of SARS-CoV-2 Mutations. Front. Microbiol., 11, 1–13.

30. Madhugiri,R., Karl,N., Petersen,D., Lamkiewicz,K. and Fricke,M. (2020) Since January 2020 Elsevier has created a COVID-19 resource centre with free information in English and Mandarin on the novel coronavirus COVID-19. The COVID-19 resource centre is hosted on Elsevier Connect , the company’s public news and information.

31. Popenda,M., Szachniuk,M., Antczak,M., Purzycka,K.J., Lukasiak,P., Bartol,N., Blazewicz,J. and Adamiak,R.W. (2012) Automated 3D structure composition for large RNAs. Nucleic Acids Res., 40, 1–12.

32. Miladi,M., Raden,M. and Diederichs,S. (2020) MutaRNA: analysis and visualization of mutation-induced changes in RNA structure. 48, 287–291.

33. Lu,X.J. and Olson,W.K. (2003) 3DNA: A software package for the analysis, rebuilding and visualization of three-dimensional nucleic acid structures. Nucleic Acids Res., 31, 5108–5121.

34. Panwar,B. and Raghava,G.P.S. (2015) Identification of protein-interacting nucleotides in a RNA sequence using composition profile of tri-nucleotides. Genomics, 105, 197–203.

35. Dijk,M. Van and Visscher,K.M. (2013) Solvated protein – DNA docking using HADDOCK. 10.1007/s10858-013-9734-x.

36. Van Dijk,M., Van Dijk,A.D.J., Hsu,V., Rolf,B. and Bonvin,A.M.J.J. (2006) Information-driven protein-DNA docking using HADDOCK: It is a matter of flexibility. Nucleic Acids Res., 34, 3317–3325.

37. Schneidman-duhovny,D., Inbar,Y., Nussinov,R. and Wolfson,H.J. (2005) PatchDock and SymmDock: servers for rigid and symmetric docking. 33, 363–367.

38. Tuszynska,I., Magnus,M., Jonak,K., Dawson,W. and Bujnicki,J.M. (2015) NPDock: A web server for protein-nucleic acid docking. Nucleic Acids Res., 43, W425–W430.

39. Yan,Y., Zhang,D., Zhou,P., Li,B. and Huang,S. (2017) HDOCK: a web server for protein – protein and protein – DNA / RNA docking based on a hybrid strategy. 45, 365–373.

40. Bowers,K.J., Chow,E., Xu,H., Dror,R.O., Eastwood,M.P., Gregersen,B.A., Klepeis,J.L., Kolossvary,I., Moraes,M.A., Sacerdoti,F.D., et al. (2006) Scalable Algorithms for Molecular Dynamics Simulations on Commodity Clusters.

41. Beard,H., Cholleti,A., Pearlman,D., Sherman,W. and Loving,K.A. (2013) Applying Physics-Based Scoring to Calculate Free Energies of Binding for Single Amino Acid Mutations in Protein-Protein Complexes. 8, 1–11.

42. Grant,B.J., Skjærven,L. and Yao,X.Q. (2020) The Bio3D packages for structural bioinformatics. Protein Sci., 10.1002/pro.3923.

43. Grant,B.J., Rodrigues,A.P.C., ElSawy,K.M., McCammon,J.A. and Caves,L.S.D. (2006) Bio3d: An R package for the comparative analysis of protein structures. Bioinformatics, 22, 2695–2696.

44. Skjaerven,L., Yao,X.Q., Scarabelli,G. and Grant,B.J. (2014) Integrating protein structural dynamics and evolutionary analysis with Bio3D. BMC Bioinformatics, 15, 1–11.

45. Skjærven,L., Jariwala,S., Yao,X.Q. and Grant,B.J. (2016) Online interactive analysis of protein structure ensembles with Bio3D-web. Bioinformatics, 32, 3510–3512.

46. Shen,X. and Masters,P.S. (2000) Evaluation of the role of heterogeneous nuclear ribonucleoprotein A1 as a host factor in murine coronavirus discontinuous transcription and genome replication.

47. Luo,H., Chen,Q., Chen,J., Chen,K., Shen,X. and Jiang,H. (2005) The nucleocapsid protein of SARS coronavirus has a high binding affinity to the human cellular heterogeneous nuclear ribonucleoprotein A1. FEBS Lett., 579, 2623–2628.

48. Elbe,S. and Buckland-Merrett,G. (2017) Data, disease and diplomacy: GISAID’s innovative contribution to global health. Glob. Challenges, 1, 33–46.

49. Kobori,T., Iwamoto,S., Takeyasu,K. and Ohtani,T. (2007) Biopolymers Volume 85 / Number 4 295. Biopolymers, 85, 392–406.

50. Kelsey C. Martin Mhatre V. Ho,J.-A.L. (2012) 基因的改变NIH Public Access. Bone, 23, 1–7.

51. Sponer,J., Bussi,G., Krepl,M., Banas,P., Bottaro,S., Cunha,R.A., Gil-Ley,A., Pinamonti,G., Poblete,S., Jurečka,P., et al. (2018) RNA structural dynamics as captured by molecular simulations: A comprehensive overview. Chem. Rev., 118, 4177–4338.

52. Humphrey,W., Dalke,A. and Schulten,K. (1996) VMD: Visual Molecular Dynamics. 7855, 33–38.

