## Supplemental Figures for "In-Silico analysis reveals lower transcription efficiency of C241T variant of SARS-CoV-2 with host replication factors MADP1 and HNRNP-1"

### Images

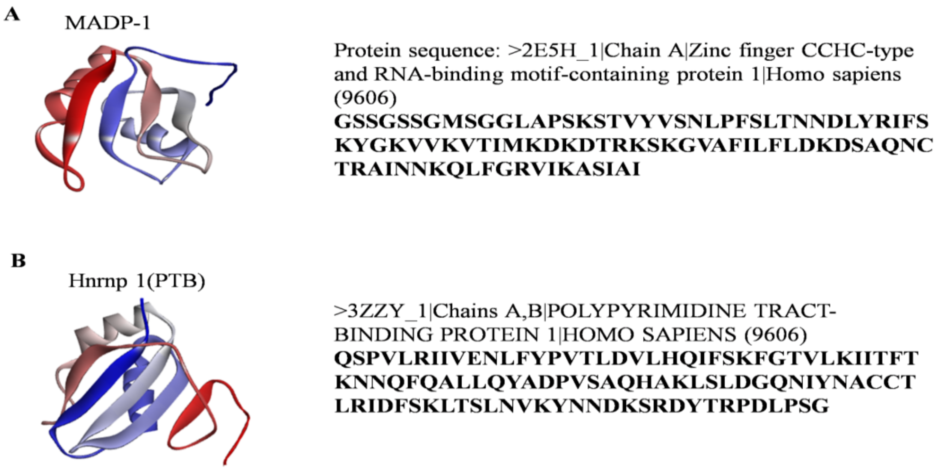


**Supplementary Figure S1:** MADP1 (**1A)** andhnRNP1 (**1B**) are used in this study and considered as a host transcription factor.

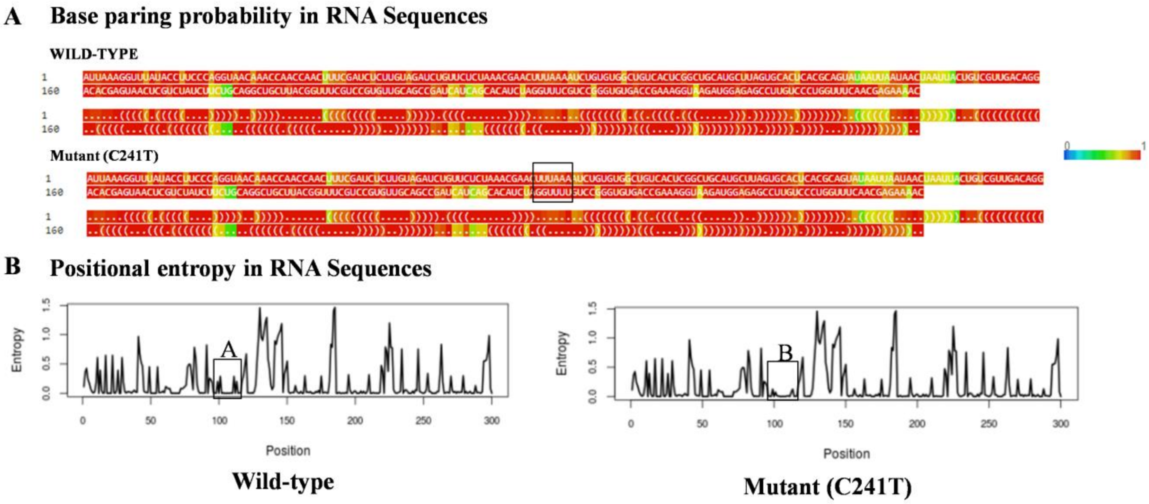


**Supplementary Figure S2:** RNA fold results for both sequences and change in base pairing probabilities

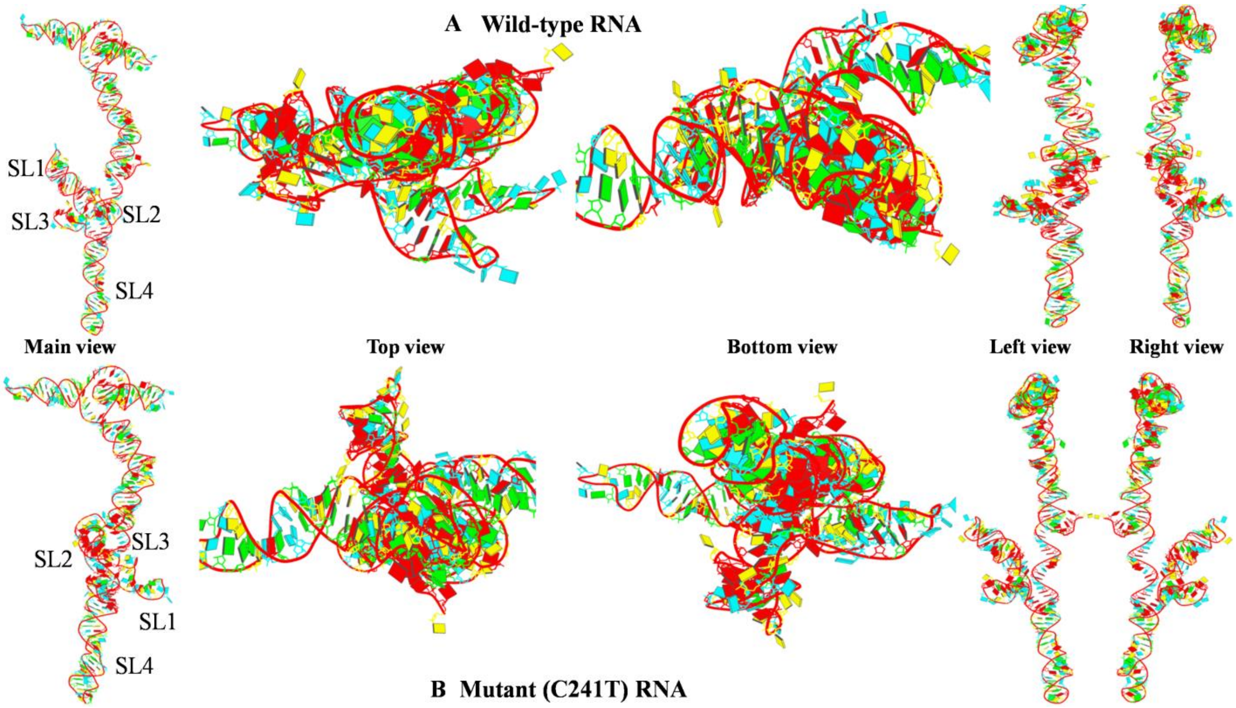


**Supplementary FigureS3:** RNA 3D structure geometry is shown. RNA pdb file generated in X3-DNA DSSR shows differences within the folding of SL1, SL2, SL3 and SL4, which can affect the protein binding within RNA.**3A:** Wild-type RNA **3B:** Mutant RNA

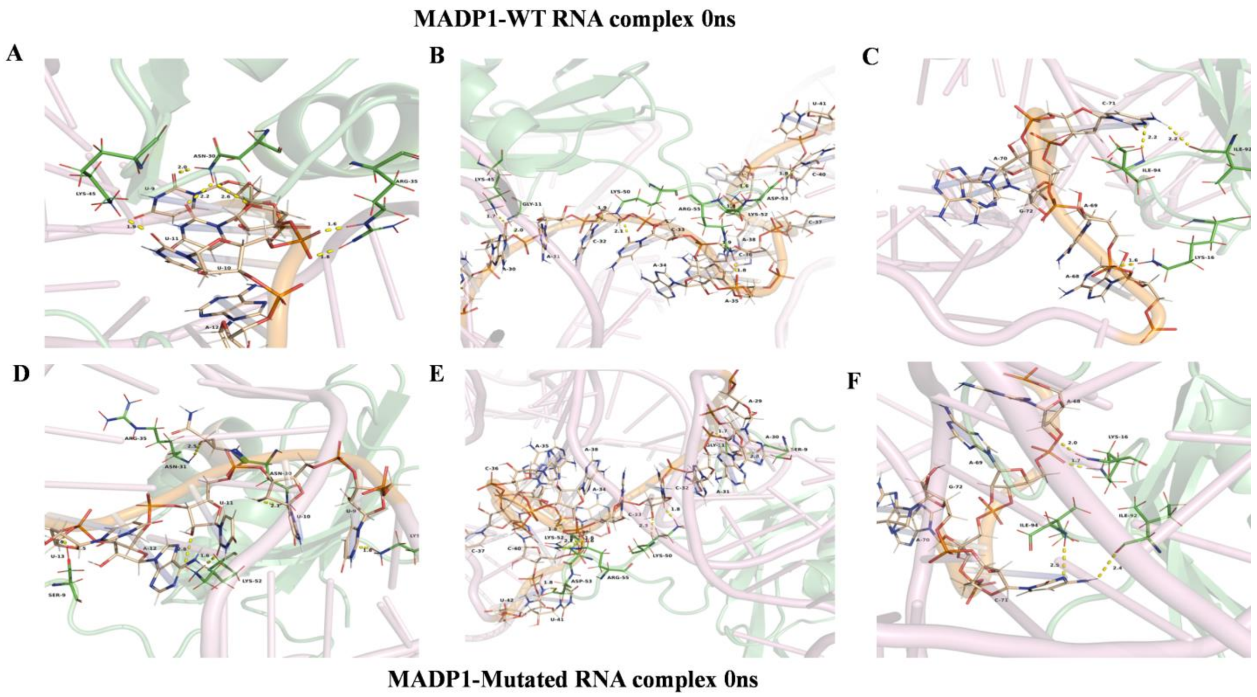


**Supplementary Figure S4:** Hydrogen bonds formation during 0ns time interval in MADP1-WT complex and mutant complex in SL1, SL2 and SL3 region.**4A & 4B**: MAPD1-WT RNA complex having pivotal interactions at SL1 region. **4C:** MAPD1-WT RNA complex having unique interactions in SL3. **4D & 4E**: MAPD1-Mutated RNA complex having pivotal interactions at SL1 region. **4F:** MAPD1-Muatated RNA complex having unique interactions in SL3.

MADP1-Wildtype Complex at 30 ns

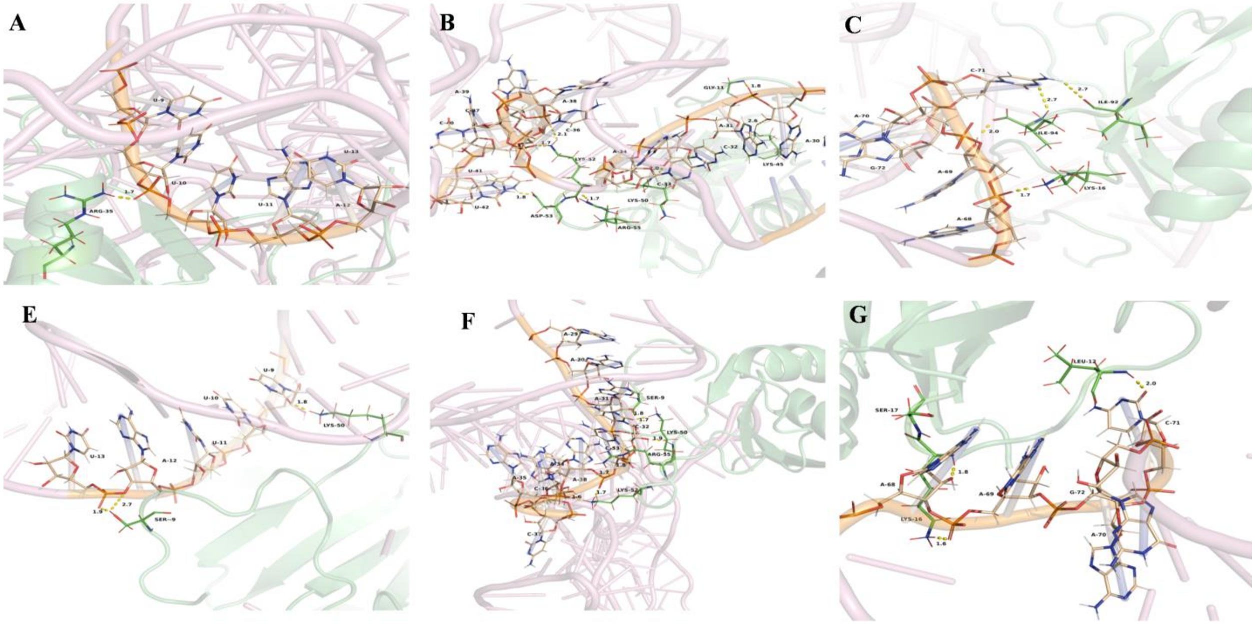


MADP1-Mutated Complex at 30 ns

**Supplementary Figure S5:** Hydrogen bonds formation during 30ns time interval lin MADP1-WT and Mutant complex in SL1, SL2 and SL3 region.**5A & 5B**: MAPD1-WT RNA complex having pivotal interactions at SL1 region. **5C:** MAPD1-WT RNA complex having unique interactions in SL3. **5D & 5E**: MAPD1-Mutated RNA complex having pivotal interactions at SL1 region. **5F:** MAPD1-Muatated RNA complex having unique interactions in SL3.

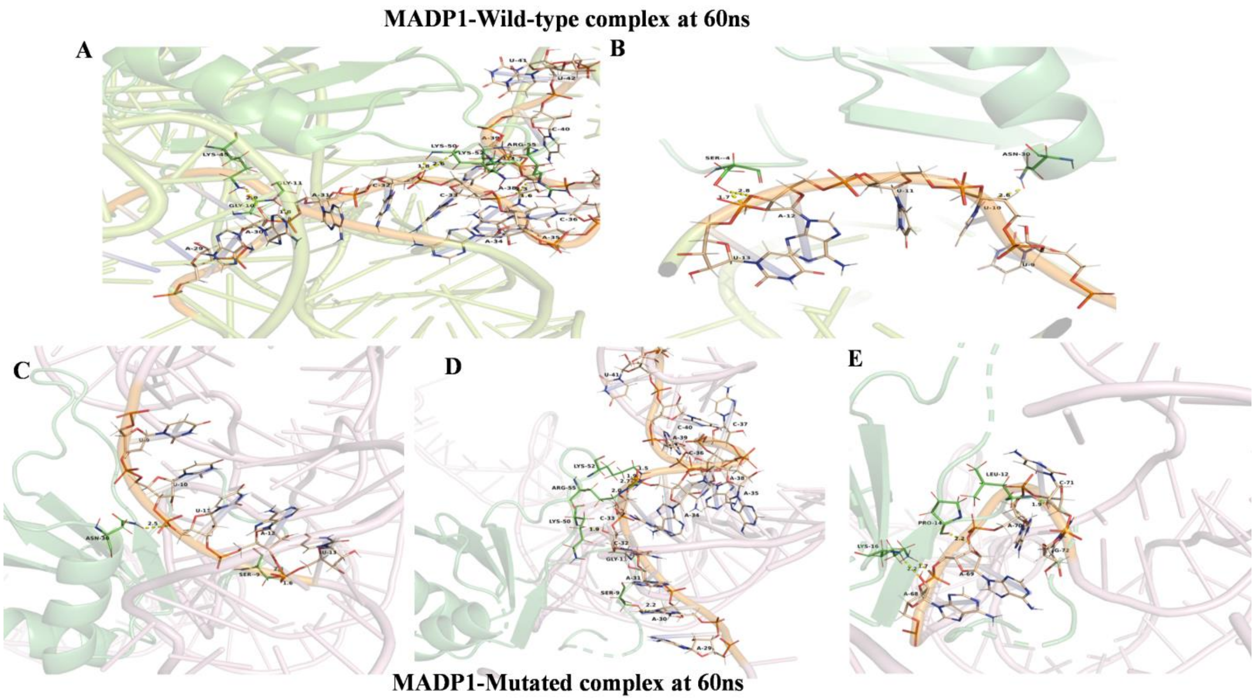


**Supplementary Figure S6:** Hydrogen bonds formation during 60ns time intervalin MADP1-WT and Mutant complex in SL1, SL2 and SL3 region.**6A & 6B**: MAPD1-WT RNA complex having pivotal interactions at SL1 region. **6C & 6D**: MAPD1-Mutated RNA complex having pivotal interactions at SL1 region. **6E:** MAPD1-Muatated RNA complex having unique interactions in SL3.

**MADP1-Wild-type complex at 100ns**

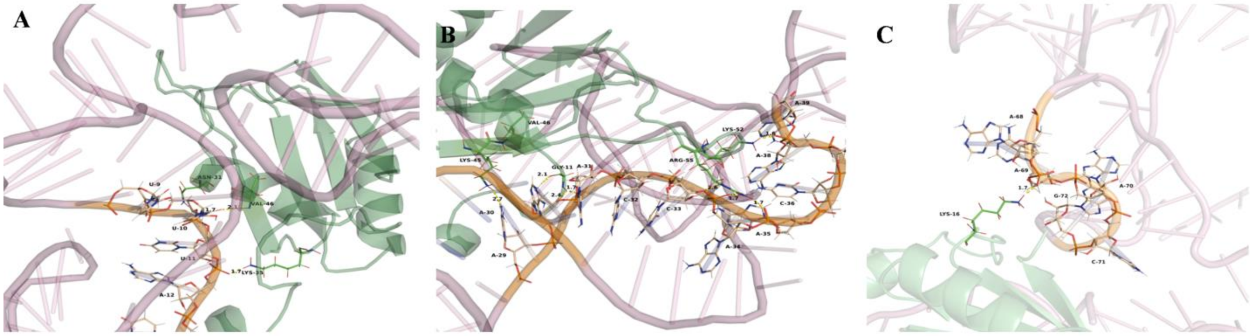


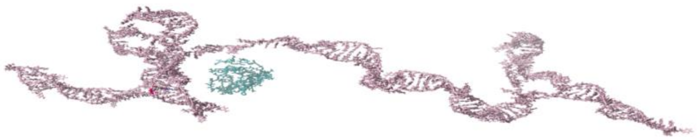

**MADP1-Mutated complex at 100ns**

**Supplementary Figure S7:** Hydrogen bonds formation during 1000ns time intervalin MADP1-WT and the Mutant complex in SL1, SL2 and SL3 region. **7A & 7B**: MAPD1-WT RNA complex having pivotal interactions at SL1 region. **7C:** MAPD1-WT RNA complex having unique interactions in SL3. **Mutated complex**: Dissociation of protein NRA was observed

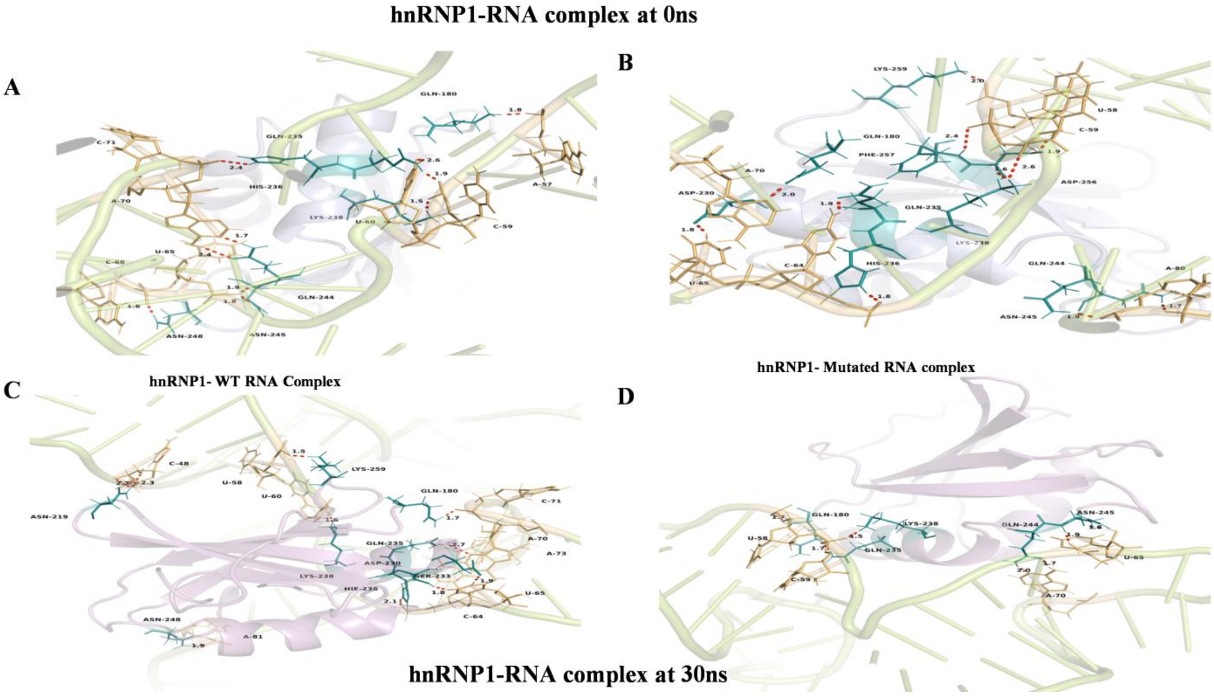


**Supplementary Figure S8:** Hydrogen bonds formation during 0ns and 30ns time interval in hnRNP1-WT and Mutant complex in SL3 region. **8A**: hnRNP1-WT RNA complex having pivotal interactions at SL3 region containing TLR-S at 0ns. **8C:** hnRNP1-Mutated RNA complex having interactions in SL3 region containing TLR-S at 0ns. **8A**: hnRNP1-WT RNA complex having pivotal interactions at SL3 region containing TLR-S at 30ns. **8C:** hnRNP1-Mutated RNA complex having interactions in SL3 region containing TLR-S at 30ns.

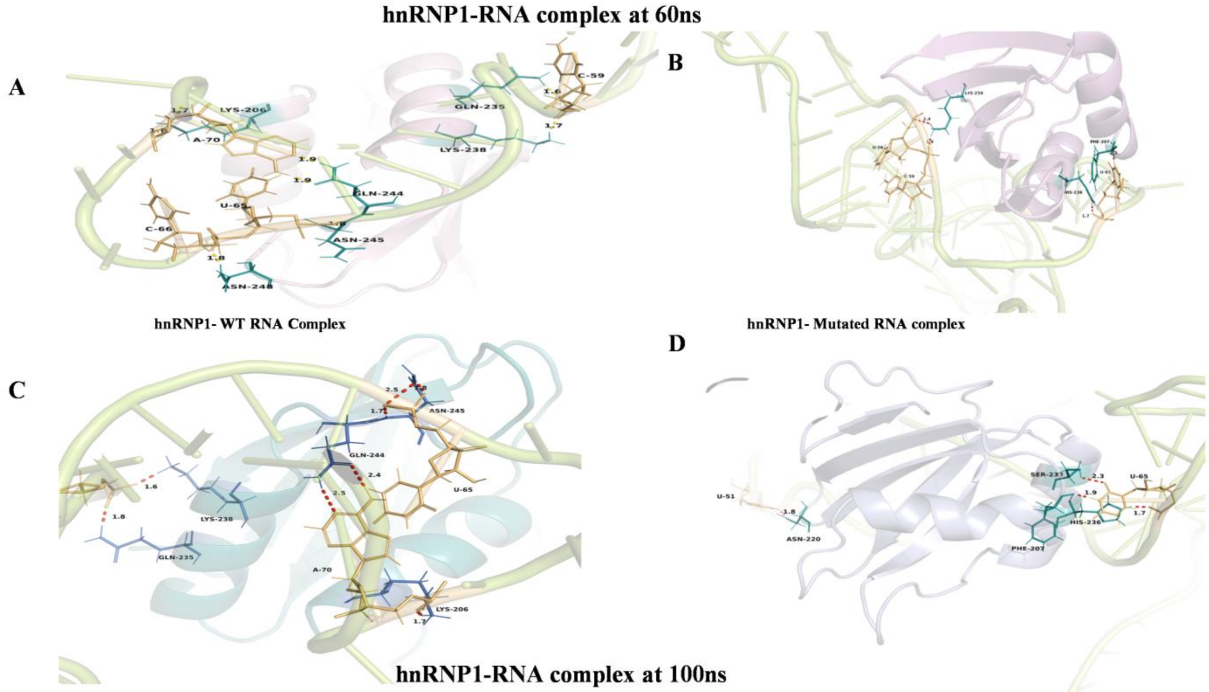


**Supplementary FigureS9:** Hydrogen bonds formation during 60ns and 100nstime interval in hnRNP1-WT and Mutant complex in SL3 region. Hydrogen bonds formation during 0ns and 30ns time intervalin hnRNP1-WT and Mutant complex in SL3 region. **9A**: hnRNP1-WT RNA complex having pivotal interactions at SL3 region containing TLR-S at 60ns. **9C:**hnRNP1-Mutated RNA complex having interactions in SL3 region containing TLR-S at 60ns. **9A**: hnRNP1-WT RNA complex having pivotal interactions at SL3 region containing TLR-S at 100ns. **9C:**hnRNP1-Mutated RNA complex having interactions in SL3 region containing TLR-S at 100ns.

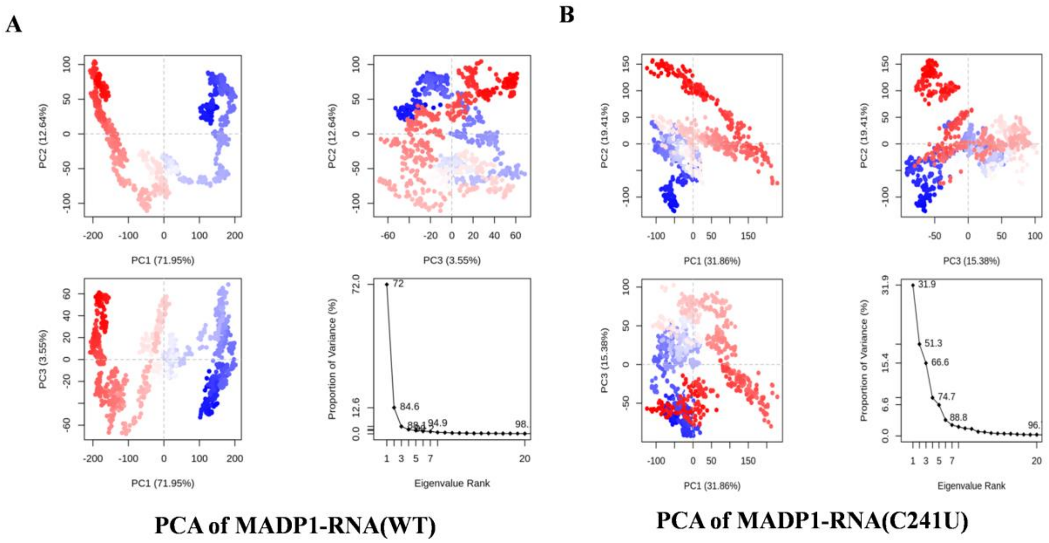


**Supplementary figure S10:** Principle component analysis for MADP1-RNA complexes. Each point represents its conformation of the protein-RNA complex throughout the X and Y axis. The spread of blue and red color dots described the degree of conformational changes in the simulation. Red color denotes compact and stable system while blue color indicates scattered and unstable system **10A:** PCA of MADP-WT RNA complex. **10B:** PCA of MADP-mutated RNA complex

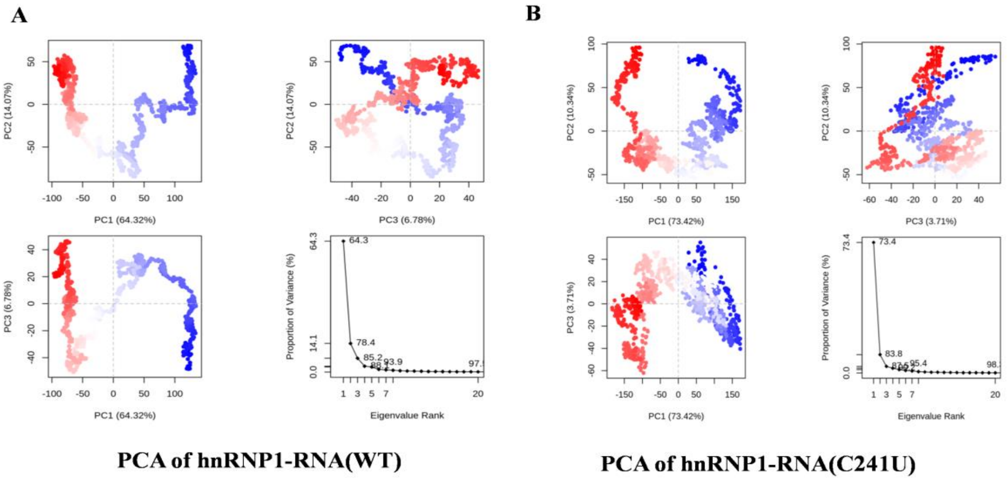


**Supplementary figure S11:** Principle component analysis for hnRNP1-RNA complexes.Each point represents its conformation of the protein-RNA complex throughout the X and Y axis. The spread of blue and red color dots described the degree of conformational changes in the simulation. Red color denotes compact and stable system while blue color indicates scattered and unstable system **11A:** PCA of hnRNP1-WT RNA complex. **11B:**PCA of hnRNP1-mutated RNA complex

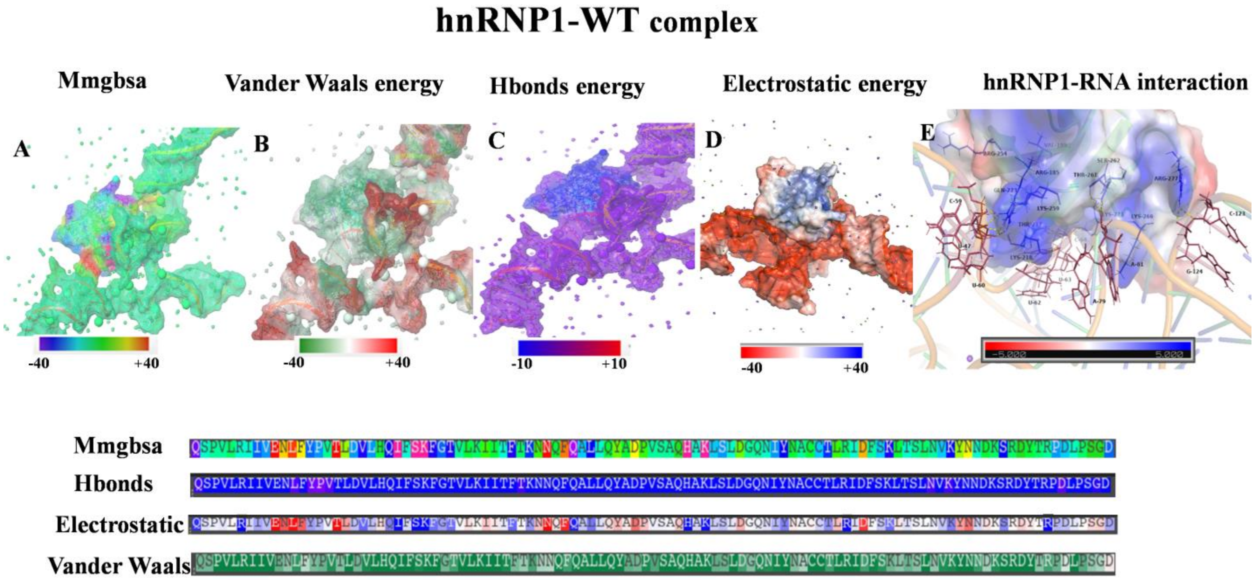


**Supplementary Figure S12:** Contribution of Major energy components involved in formation of hnRNP1-Wild-type RNA complex**.** Energy components contribution was done in energy visualization package in maestro by entering cut off values of -40 to +40 kcal/mole for MMGBSA, electrostatic potential, and Vander Waals energy. -10 to +10 cut-off values for Hydrogen bond energy.**12A:** MMGBSA for complex. **12B:** Vander Waals energy **12C:** Hbonds energy **12D:** Electrostatic energy and **12E: hnRNP1**-WT RNA interaction in terms of electrostatic energy, as RNA was showing negative electrostatic potential due to the phosphate backbone it was not shown for better image of interaction (for whole electrostatic interaction figure **12D** can be referred)

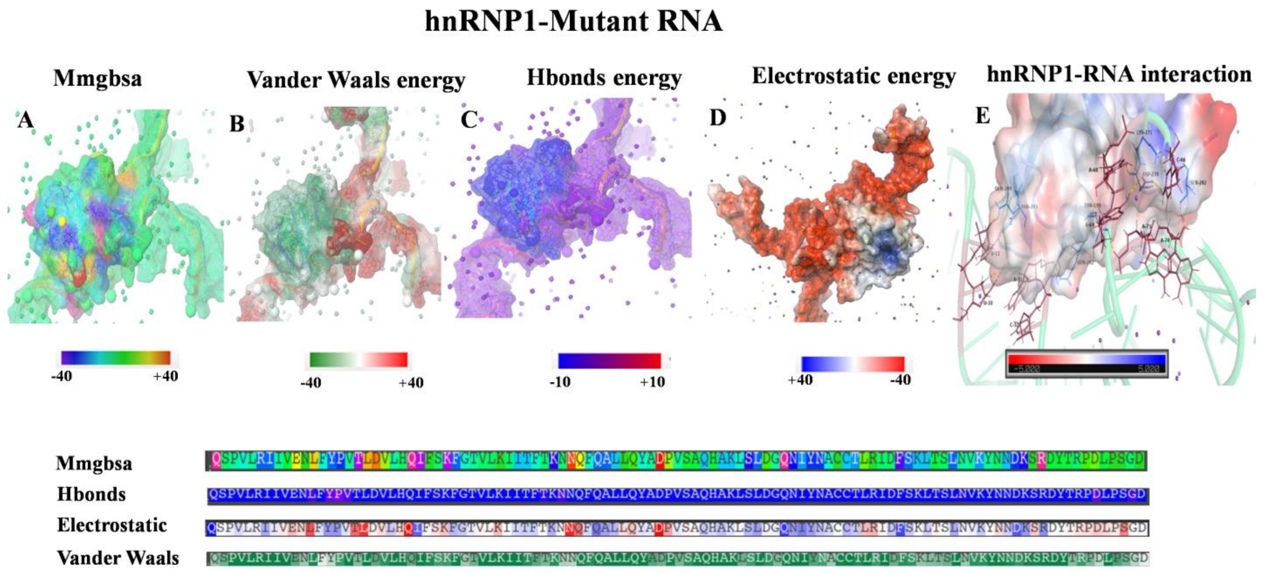


**Supplementary figure S13:** Contribution of Major energy components involved in formation of hnRNP1-Wild-type RNA complex**.** Energy components contribution was done in energy visualization package in maestro by entering cut off values of -40 to +40 kcal/mole for MMGBSA, electrostatic potential, and Vander Waals energy. -10 to +10 cut-off values for Hydrogen bond energy.**13A:** MMGBSA for complex. **13B:** Vander Waals energy **13C:** Hbonds energy **13D:** Electrostatic energy and **13E:** hnRNP1-WT RNA interaction in terms of electrostatic energy, as RNA was showing negative electrostatic potential due to the phosphate backbone it was not shown for better image of interaction (for whole electrostatic interaction figure **13E** can be referred)

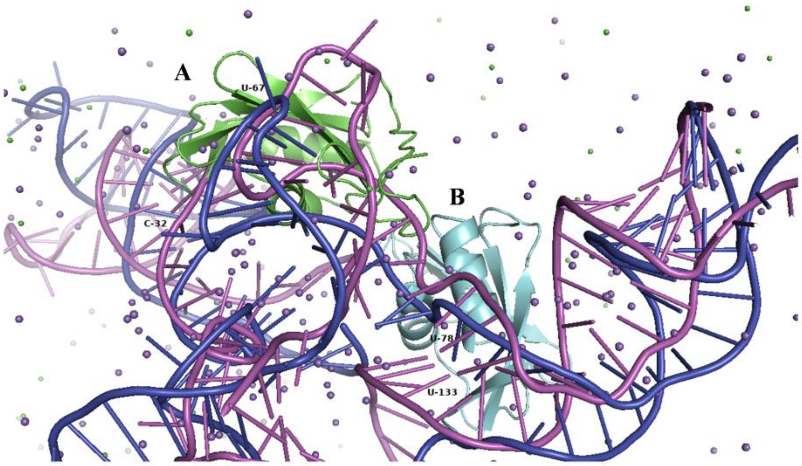


**Minimum energy structure generated from MMGBSA method for hnRNP1-RNA complexes**

**Supplementary figure S14: MMGBSA minimum energy structures for hnRNP1.**

**A:** A denotes wild-type hnRNP1 where it is bound near to SL1 and SL3 and **B:** denotes Mutant hnRNP1 where it bound near to SL3 and SL4

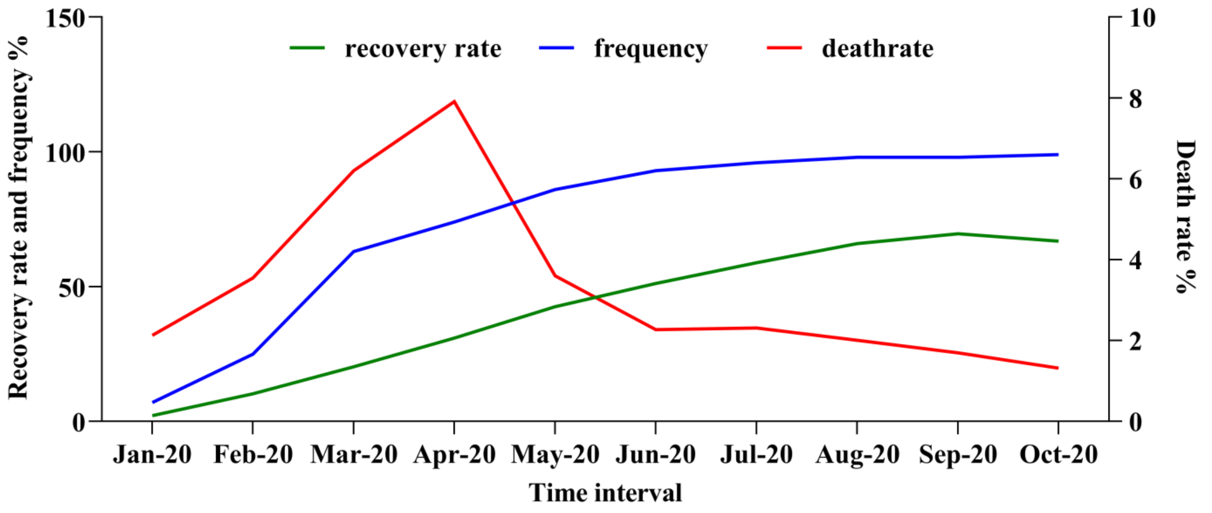


**Supplementary figure: S15 Epidemiological statistics:** Death rate and recovery rate was obtained using WHO covid-19 dashboard. Mutation frequency is obtained through NextStrain

**Supplementary Table S1:** RNA sequences taken for study

| **Types** | **RNA nucleotide sequence** | **RNA sequence in dot bracket form** |
|---|---|---|
| **Wild-type** | AUUAAAGGUUUAUACCUUCCCAGGUAAC AAACCAACCAACUUUCGAUCUCUUGUAG AUCUGUUCUCUAAACGAACUUUAAAAUC UGUGUGGCUGUCACUCGGCUGCAUGCUU AGUGCACUCACGCAGUAUAAUUAAUAAC UAAUUACUGUCGUUGACAGGACACGAGU AACUCGUCUAUCUUCUGCAGGCUGCUUAC GGUUUCGUCCGUGUUGCAGCCGAUCAUC AGCACAUCUAGGUUUCGUCCGGGUGUGA CCGAAAGGUAAGAUGGAGAGCCUUGUCC CUGGUUUCAACGAGAAAAC | ......(((((.(((((....)))))..)))))...........((( ((.....))))).((((.......))))........(((((((.... .((((.(((.....))).))))....)))))))............... .......(((((((((((..(((((...(((.((((((((..... ((((((.(((((......)))))..))))))..........((((( ((.((......)))))))))(((....))))))))))))))).)) ))).))))...)))))))...... |
| **Mutant(C241U)** | AUUAAAGGUUUAUACCUUCCCAGGUAAC AAACCAACCAACUUUCGAUCUCUUGUAG AUCUGUUCUCUAAACGAACUUUAAAAUC UGUGUGGCUGUCACUCGGCUGCAUGCUU AGUGCACUCACGCAGUAUAAUUAAUAAC UAAUUACUGUCGUUGACAGGACACGAGU AACUCGUCUAUCUUCUGCAGGCUGCUUAC GGUUUCGUCCGUGUUGCAGCCGAUCAUC AGCACAUCUAGGUUUUGUCCGGGUGUGA CCGAAAGGUAAGAUGGAGAGCCUUGUCC CUGGUUUCAACGAGAAAAC | ......(((((.(((((....)))))..)))))...........((( ((.....))))).((((.......))))........(((((((.... .((((.(((.....)).)))))....)))))))............... .......(((((((((((..(((((...(((.((((((((..... ((((((.(((((......)))))..))))))..........((((( ((.((......)))))))))(((....))))))))))))))).)) ))).))))...)))))))...... |

**Supplementary table S2.**Hydrogen bond occupancy within MADP-RNA complexes with respect to SL1 and SL2

| **Sr.N0** | **MADP1-Wild-type** | | | **MADP1-Mutant (C241U)** | | |
|---|---|---|---|---|---|---|
| 1 | MET-8-Side-N | U11-Side-OP1' | 87.56% | LYS39-Side-N | C72-Side-O4' | 35.36% |
| 2 | SER6-Side-OG | C21-Side-C3 | 20.06% | TYR34-Side-OH | A22-Side-OP | 41.25% |
| 3 | GLY7-Side-OG | C19-Side-O2' | 68.26% | ASN31-Side-U18 | U18-Side-O4' | 55.32% |
| 4 | SER9-Side-OG | G24-Side-O2' | 96.65% | ASP67-Side-CA | A12-Side-OP' | 35.62% |
| 5 | LYS50-Side-N’ | G8-Side-N2' | 52.50% | SER70-Side-OG | A12-Side-O4' | 43.62% |
| 6 | ARG55-Main-NH2 | G8-Side-OP1 | 20.63% | SER-9-Main-O | A12-Side-OP1 | 45.23% |
| 5 | LYS39-side-NH1 | C21-side-O3 | 53.19% | ARG55-side-NH1 | C33-side-C4’ | 55.21% |
| 6 | ARG35-Side-NH2 | C21-Side-OP2' | 85.05% | LYS52-Side-NH2 | A39-Side-O4' | 74.12% |
| 7 | ARG35-side-NH1 | U10-Side-OP1 | 60.23% | THR29-side-22A | A22-Side-OP1 | 23.41% |
| 8 | THR54-Main-NZ | U43-Side-O3' | 14.97% | LYS16-Side-NZ | A68-Side-O3' | 75.52% |
| 9 | LYS45-Side-NH | C33-Side-C5' | 88.32% | VAL46-Side-NZ | U10-Side-O4' | 64.21% |
| 10 | ILE48-Main-CD1 | G24-Side-P' | 93.99% | LYS58-Side-NH | U42-Side-O4' | 54.23% |
| 11 | ASP53-Side-OD | U42-Side-C4 | 36.33% | SER4-Side-OH | A30-Side-OP2 | 22.35% |
| 12 | LYS68-side-NH | U67-Side-C5 | 30.39% | GLY11-side-N | U12-Side-C5 | 2.36% |

**Supplementary table S3:**Hydrogen bond occupancy within hnRNP1-RNA complexes with respect to SL3

| Sr.NO | HnRNP1-Wild-type | | | HnRNP1-Mutant (C241U) | | |
|---|---|---|---|---|---|---|
| 1 | GLN180-Side-N | G56-Side-OP1’ | 87.56% | GLN180-Side-N | A70-Side-O3' | 84.36% |
| 2 | GLN235-Side-OE | U58-Side-O4’ | 80.06% | GLN235-Side-OE | U64-Side-N3 | 61.25% |
| 3 | LYS238-Side-NZ | U58-Side-OP' | 68.26% | LYS238-Side-NZ | U59-Side-O2' | 75.32% |
| 4 | ASN245-Side-NE | C64-Side-O2' | 96.65% | ASN245-Side-NE | A80-Side-O2' | 85.62% |
| 5 | LEU253-Side-N | U65-Side-C4' | 52.50% | LEU235-Side-N | U63-Side-O4' | 43.62% |
| 6 | ASN248-Main-O | U65-Side-OP1 | 20.63% | ASN248-Main-O | C64-Side-OP1 | 38.23% |
| 5 | GLN244-side-NH1 | A70-side-O3’ | 53.19% | GLN244-side-NH1 | A79-side-OP | 75.21% |
| 6 | HIS236-Side-NH2 | A70-Side-OP2' | 85.05% | TYR247-Side-NH2 | U65-Side-O4' | 74.12% |
| 7 | LEU253-side-NH2 | U75-Side-OP1 | 82.23% | VAL232-side-NH2 | C66-Side-OP1 | 53.41% |
| 8 | LEU241-Main-NZ | A74-Side-O3' | 44.97% | HIS236-Main-NZ | C64-Side-O3' | 71.52% |
| 9 | GLN223-Side-OE1 | C59-Side-C5' | 88.32% | LYS38-Side-OG | C75-Side-O4' | 64.21% |
| 10 | LYS266-Side-NH1 | A81-Side-O4' | 93.99% | VAL232-Side-OG | C66-Side-O4' | 54.23% |
| 11 | ARG277-Side-NH2 | U77-Side-OP2 | 36.33% | LYS206-Side-NH1 | A68-Side-OP2 | 35.35% |
| 12 | LYS271-side-NH1 | A81-Side-C5 | 93.39% | LYS271-Side-NH2 | C66-Side-C5 | 82.36% |

**Supplementary video S1:** MADP-1 Wild-type complex having no dissociation at 100ns

**Supplementary video S2**: MADP1-Muated complex started getting dissociation after 97 ns.
